# Non-ionic CT Contrast Solutions Rapidly Alter Bovine Cartilage and Meniscus Mechanics

**DOI:** 10.1101/2019.12.13.874743

**Authors:** Eva G. Baylon, Hollis A. Crowder, Garry E. Gold, Marc E. Levenston

**Affiliations:** Department of Mechanical Engineering, Stanford University, Stanford, CA 94305; Department of Radiology, Stanford University, Stanford, CA 94305; Department of Bioengineering, Stanford University, Stanford, CA 94305; Department of Orthopaedic Surgery, Stanford University, Stanford, CA 94305

**Keywords:** contrast agent, contact modulus, cartilage, meniscus, osmolality

## Abstract

**Objective:** To evaluate effects of a common CT contrast agent (iohexol) on the mechanical behaviors of cartilage and meniscus.

**Methods:** Indentation responses of juvenile bovine cartilage and meniscus were monitored following exposure to undiluted contrast agent (100% CA), 50% CA/water, 50% CA/Phosphate Buffered Saline (PBS) or PBS alone, and during re-equilibration in PBS. The normalized peak force 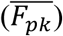, effective osmotic strain (*ε*_*osm*_), and normalized effective contact modulus 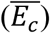 were calculated for every cycle, with time constants determined for both exposure and recovery via mono- or biexponential fits to 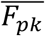.

**Results:** All cartilage CA groups exhibited long-term increases in 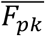 following exposure, although the hyperosmolal 100% CA and 50% CA/PBS groups showed an initial transient decrease. Meniscus presented opposing trends, with decreasing 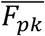 for all CA groups. Re-equilibration in PBS for 1hr after exposure to 100% CA did not produce recovery to baseline 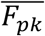 in either tissue. The recovery time for meniscus was substantially longer than that of cartilage. 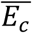 increased with CA exposure time for cartilage but decreased for meniscus, suggesting an increased effective stiffness for cartilage and decreased stiffness for meniscus. Long-term changes to *ε*_*osm*_ in both tissues were consistent with changes in 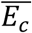.

**Conclusion:** Exposure to iohexol solutions affected joint tissues differentially, with increased cartilage stiffness, likely relating to competing hyperosmotic and hypotonic interactions with tissue fixed charges, and decreased meniscus stiffness, likely dominated by hyperosmolarity. These altered tissue mechanics could allow non-physiological deformation during ambulatory weight-bearing, resulting in an increased risk of tissue or cell damage.

## Introduction

Computed Tomography (CT) arthrography is used to assess soft tissue structures and surface lesions across a range of pathologies and degenerative conditions^1,2^, particularly in the shoulder, elbow, hip and knee, and recent studies have explored its utility to estimate glycosaminoglycan (GAG) composition in cartilage^3,4^. Conventional CT arthrograms of the knee involve intraarticular injection of an iodinated contrast agent followed by a CT scan. While magnetic resonance (MR) arthrography is more common and minimizes exposure to ionizing radiation, CT arthrography continues to be useful due to its broad availability, lower cost, and its utility in patients with contraindications to MRI, including prostheses or metal implants^5,6^. Additionally, CT contrast agents are often included in MR contrast injections for fluoroscopic guidance during MR arthrography or to allow sequential MR and CT arthrography.

Iodine is the basis for many CT contrast agents, as its high X-ray attenuation enhances visibility of blood vessels and tissues^7^. Both ionic and non-ionic contrast agents have been widely used in clinical and research settings, but non-ionic agents have become favored due to lower osmolalities and decreased physiological side effects^8,9^. Although most non-ionic, iodinated contrast agents are considered low-osmolar, their osmolalities range from ∼300-800 mOsm/kg and most are hyperosmotic relative to normal saline (308 mOsm/kg) and synovial fluid (300-400mOsm/kg)^10–12^. Importantly, these contrast agents are solubilized in water with minimal ionic excipients, giving these hyperosmolal non-ionic contrast solutions substantially lower ionic strengths than synovial fluid.

Despite the prevalence of intraarticular contrast injections, formulations for injection are based primarily on institutional preference and expected image quality rather than specific considerations of effects on articular tissues^13,14^. Protocols vary substantially and involve contrast agents undiluted or variably diluted in anesthetics or saline^3,13,15–20^. Although the full range of clinical protocols is unknown, contrast formulations reported in the recent research literature vary substantially (Fig. 1, Table S1). Ionic contrast formulations had hyperphysiologic ionic strengths and osmolalities up to 5 times that of synovial fluid, while non-ionic contrast formulations had subphysiological (in some cases negligible) ionic strengths and generally had hyperphysiologic osmolalities up to twice that of synovial fluid.

**Figure 1:**
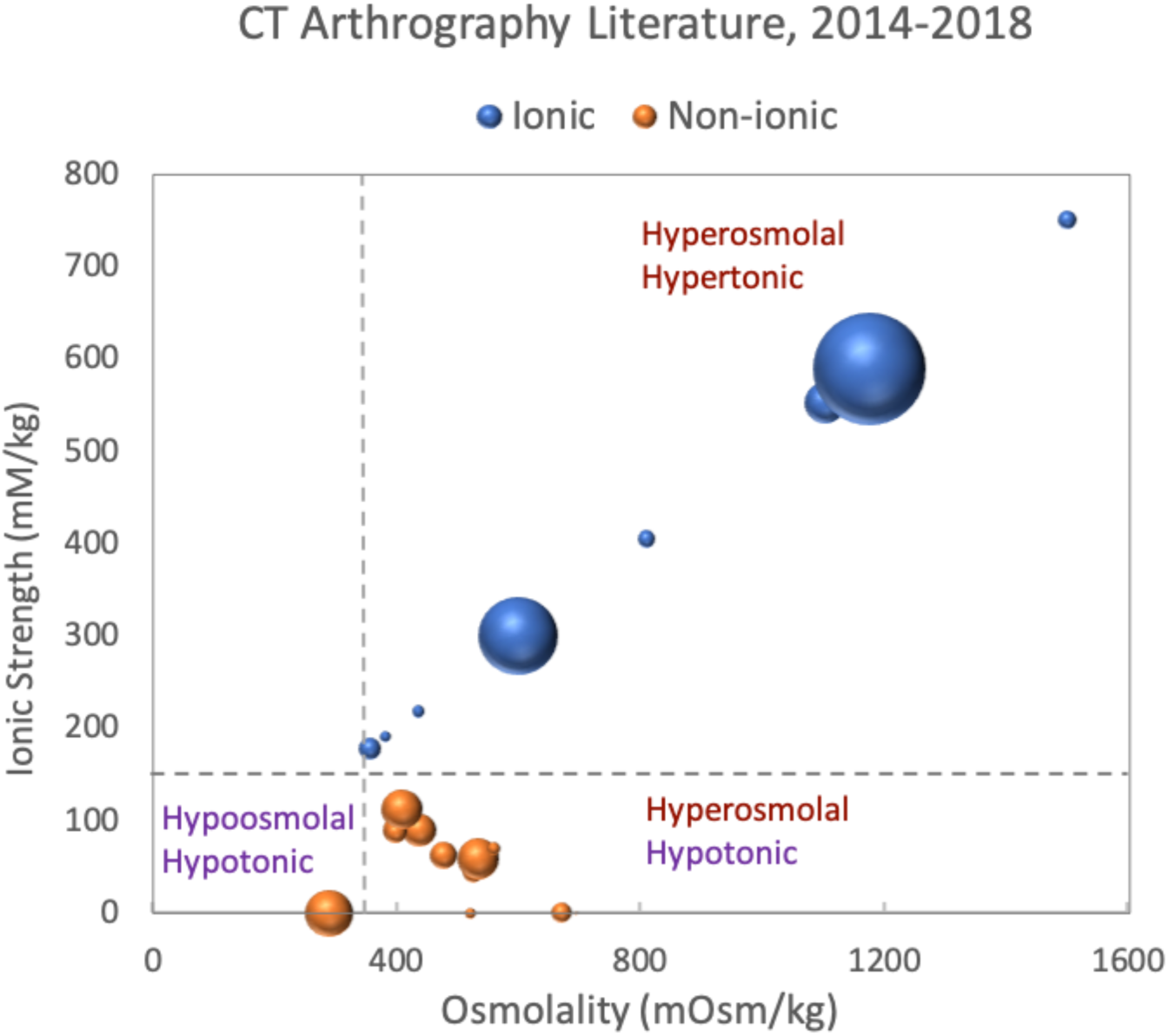
Contrast agents for CT arthrography vary substantially in both osmolality and ionic strength. Osmolalities and ionic strengths were estimated for all PubMed-indexed CT arthrography studies on human subjects published in 2014-2018 that gave sufficient information on the contrast agent and dilution (47 studies). Bubble sizes indicate the number of human subjects for a given contrast formulation, pooling multiple studies with identical formulations. Dashed lines indicate approximate values for synovial fluid. Ionic contrast solutions were generally both hyperosmolal and hypertonic with respect to synovial fluid, with the largest set of studies (largest bubble represents ∼1500 subjects) reporting use of contrast formulation with an osmolality and ionic strength 3-4 times larger than synovial fluid. Non-ionic contrast solutions were generally hyperosmolal (1-2 times larger than synovial fluid) but hypotonic with respect to synovial fluid.

These large variations in contrast solution properties are concerning given the known sensitivity of proteoglycan-rich tissues, such as cartilage, to the osmotic environment. Alterations to bath ion concentrations affect the osmotic swelling stress in cartilaginous tissues produced by interactions between the negatively charged sulfated GAGs and the ions in the interstitial fluid^21–28^. These interactions are functionally important and contribute to both the equilibrium and dynamic stiffness of these tissues. As arthrography patients often ambulate to distribute the contrast medium immediately post-injection, changes to the osmotic environment induced by contrast media could be detrimental for cartilaginous tissues. Indeed, Hirvasniemi et. al. specifically avoided subject weight-bearing based on findings that 1.5-hour immersion in a hyperosmolal, ionic contrast agent (Hexabrix 320, 600mOsm/kg) substantially decreased bovine cartilage stiffness and increased susceptibility to loading-induced cell death^3,29^. Nonetheless, osmolality alone may not be sufficient to predict the effects of a solution on cartilaginous tissues, as ionic and non-ionic solutes interact differentially with the charged tissue matrices.

Although systemic effects of iodinated contrast agents have been characterized, potential effects on the mechanical response to loading of cartilage and meniscus – tissues directly exposed to the contrast agent solutions for prolonged periods during arthrography – are virtually unknown. This study examined transient mechanical effects on cartilage and meniscus tissues of exposure to and removal of various dilutions of iohexol, a common low-osmolar, non-ionic, iodinated contrast agent.

## Materials and Methods

### Materials

Juvenile (∼2-week old) bovine stifles were from San Jose Valley Veal and Beef (Santa Clara, CA). Biopsy punches were from Integra Miltex (York, PA). Protease Inhibitor Cocktail Set I and phosphate buffered saline (PBS) tablets (140mM NaCl, 10mM phosphate buffer, 3mM KCl reconstituted in 1L of deionized H_2_O) were from Calbiochem (San Diego, CA). Omnipaque was from GE Healthcare (Chicago, IL) and contained either 350mg/mL (Omnipaque 350, 844 mOsm/kg, 541mOsm/L) or 300mg/mL (Omnipaque 300, 672mOsm/kg, 465mOsm/L) of iodine in the form of iohexol, 1.21mg/mL tromethamine, and 0.1mg/mL edetate calcium disodium, with a reported pH between 6.8-7.7^30^.

### Sample preparation

Full thickness cores were harvested with an 8mm biopsy punch from the femoral condyle of three juvenile bovine stifles (cartilage) and the cranial surfaces of medial and lateral menisci of six juvenile bovine stifles (meniscus). Cores were trimmed to 2mm from the intact surface and stored in 1X PBS with protease inhibitors at −20°C until testing. Samples were thawed at 37°C for 30 minutes and trimmed to 6mm diameter immediately before testing. Samples were randomly distributed across treatment groups.

### Protocol development

A repeated indentation testing protocol was developed to monitor time-dependent effects of the bath environment on tissue mechanical behaviors with minimal transport inhibition, while avoiding tissue damage due to repeated testing. Tissue samples were placed within a rubber confining ring (Fig. 2A) in 5mL of 1X PBS and a compressive tare load of −0.02N was applied to determine the reference thickness. The articular surface was cyclically indented using an Instron 5944 microtester (Instron, Norwood, MA) equipped with a 10N load cell and an aluminum indenter with a hemispherical 1mm diameter tip. Each cycle consisted of indentation by 15% of the sample reference thickness at 0.002mm/s, unloading at 0.002mm/s to the reference thickness and either a 3min (for cartilage) or 15min (for meniscus) hold period (Fig. 2B, 2C). The different hold periods were determined from preliminary tests in 1X PBS as sufficient to allow full recovery of each tissue with no evidence of damage from the cyclic indentations over several hours. To validate that this protocol captured expected trends with changes in the bath environment^31^, cartilage and meniscus samples (n=3/condition) were monitored during equilibration in hypoosmolal 0.1X PBS (swelling), hyperosmolal 10X PBS (deswelling) and isoosmolal 1X PBS (control).

**Figure 2:**
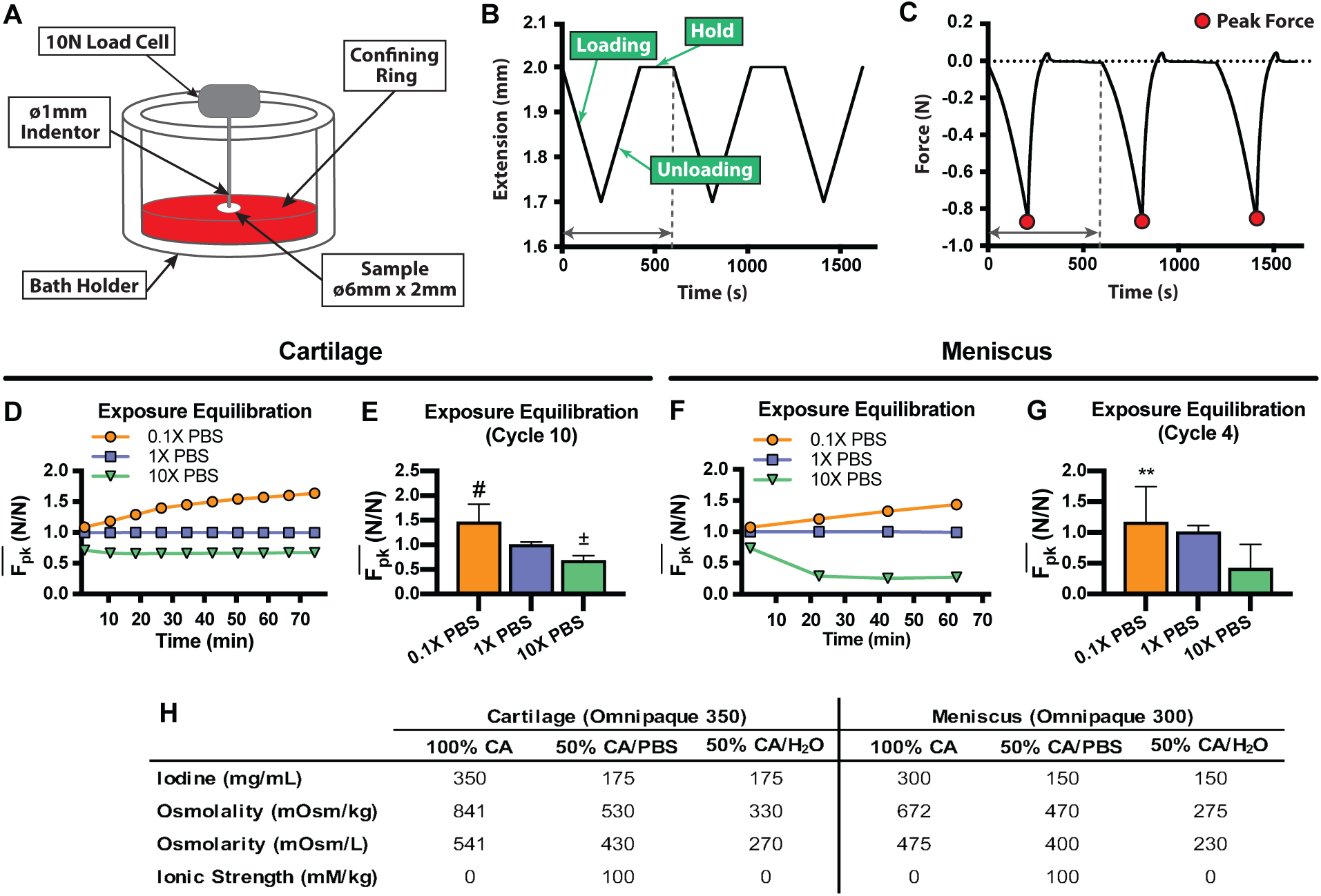
Indentation test set-up and protocol validation. (A) The indentation test set-up consisted of a thin ø1mm hemispherical indenter used to apply the prescribed strain to a ø6mm sample semi-confined in a rubber ring. (B) Each indentation cycle comprised a loading step to 15% compressive strain at a ramp rate of 0.002mm/s followed by an unloading step at the same rate and a hold period of 3 min for cartilage or 15 min for meniscus specimens; (C) the peak force for every cycle was monitored to assess changes in the mechanical response as samples were exposed to different bath solutions. The indentation test set-up was validated to assess transient changes in swelling stress by equilibrating samples in different concentrations of PBS following baseline indentation in 1X PBS (swelling: 0.1X PBS, deswelling: 10X PBS, control: 1X PBS) in (D-E) cartilage and (F-G) meniscus. As expected, normalized peak force values for samples equilibrated in 1X PBS did not change with indentation cycle, demonstrating that force changes are a result of bath equilibration and not the indentation protocol. Both (D) cartilage and (H) meniscus samples equilibrated in 0.1X PBS had increased normalized peak forces while exposure to 10X PBS resulted in decreases to the normalized peak force for both tissues. Normalized peak force at the end of cycle 10 for cartilage was significantly greater for 0.1X PBS and significantly lower for 10X PBS compared to the 1X PBS control (E). For meniscus, normalized peak force at the end of cycle 4 was significantly greater for 0.1X PBS than 1X PBS, with no significant differences detected between 10X PBS and 1X PBS (I). (H) Characteristics of contrast solutions used for cartilage and meniscus. Osmolalities, osmolarities and ionic strengths for contrast dilutions were estimated based on published physical properties of contrast agents and PBS.

### Contrast agent dilution and visual assessment of tissue effects

To examine effects of a range of clinically relevant conditions involving different osmolalities and ionic strengths, we used contrast agent as supplied (100% CA), diluted 50% v/v in PBS (50% CA/PBS), diluted 50% v/v in deionized water (50% CA/H_2_O), and 1X PBS as a control. Although clinical protocols typically use normal saline, PBS was chosen as an osmotically similar (280-310 mOsm/kg vs. 308 mOsm/kg) diluent for consistency with earlier laboratory studies. The 50% CA/H_2_O group was included to understand the effect of ions introduced by the dilution in PBS and to simulate the response to a lower osmolality, but still non-ionic contrast agent solution. Due to availability at the time of testing, cartilage was tested with Omnipaque 350 while meniscus was tested with Omnipaque 300. Published and estimated properties of the various contrast dilutions are summarized in Fig. 2H.

To visually assess any obvious changes resulting from contrast solution exposure, samples were equilibrated at room temperature in 1mL of either 100% CA, 50% CA/PBS, 50% CA/H_2_O, or 1X PBS. Photos were taken after 1hr.

### Indentation testing

Explants (n=5/condition) were subjected to the repeated indentation regime described above. Each change in bath fluid was preceded by emptying the bath and rinsing twice with the new solution. After three baseline cycles in 1X PBS, the bath was replaced by 5mL of either 100% CA, 50% CA/PBS, 50% CA/H_2_O or 1X PBS. To keep exposure times roughly comparable, cartilage samples were indented for 10 cycles while meniscus samples were indented for 4 cycles. The bath was then replaced with 1X PBS and recovery was monitored for an additional 10 (cartilage) or 4 (meniscus) cycles. Additional meniscus explants (n=3/condition) were exposed to 100% CA or 1X PBS as described above but were allowed to recover for 25 cycles in 1X PBS to monitor long-term recovery.

### Data analysis

The peak force (*F*_*pk*_) value for every cycle was normalized by *F*_*pk*_ at cycle 3, the last cycle tested in the baseline 1X PBS bath (*F*_*pk_PBS*_):

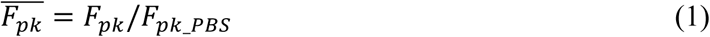

The equilibration time was characterized by fitting the normalized peak force data 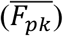 during equilibration to contrast solutions using a biexponential model:

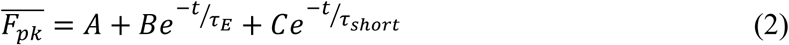

where the equilibration time constant *τ*_*E*_ describes the time scale for exposure equilibration, while *τ*_*short*_ denotes the time scale for an additional short-term response exhibited by cartilage groups (100% CA and 50% CA/PBS). Positive/negative values of the coefficients *B* and *C* indicate long-term (for B) or short-term (for C) increases/decreases to 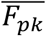. As meniscus groups exhibited a simpler response, samples from these groups were instead fitted with a monoexponential model:

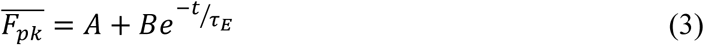

The recovery time was characterized by fitting the normalized peak force data during recovery using a monoexponential model:

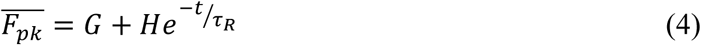

where the recovery time constant *τ*_*R*_ describes the time scale for recovery equilibration and positive/negative values of the coefficient *H* indicate long-term increases/decreases in the 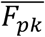 during re-equilibration in 1X PBS. As meniscus explants did not exhibit exponential behaviors during recovery, the recovery exponential fit was not performed for these groups. For 100% CA groups only, the recovery response of cartilage and meniscus was assessed by comparing the actual time values needed to reach 50% recovery (t_50%_).

Changes in tissue thickness could not be consistently determined from the force-deflection curves because samples that deswelled (contracted) developed a gap between the indenter and tissue surface while samples that swelled maintained contact. As an alternative, we determined for each cycle *d*_0.1*PF*_, the compressive displacement at which the force reached 10% of the cycle’s peak force as an estimate of whether and to what extent a sample changed dimension. An effective osmotic strain (*ε*_*osm*_) relative to baseline was calculated as:

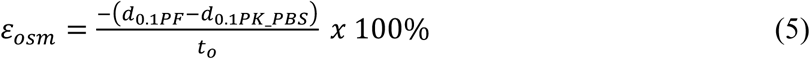

where *d*_0.1*PK_PBS*_ is the displacement at which the last baseline cycle reached 10% of its peak force and *t*_*o*_ is the reference tissue thickness. Swelling results in smaller displacement values and positive values of *ε*_*osm*_ while deswelling results in negative values of *ε*_*osm*_.

### Contact modulus

The loading portions of each indentation cycle (Fig. S1, S2) were fitted with an analytical solution to the Hertzian contact model for biphasic materials^32^ to determine an effective contact modulus (*E*_*c*_). *E*_*c*_ reflects combined fluid and solid contributions to the indentation stiffness that depends on intrinsic material properties as well as experimental conditions, such as indentation rate and probe radius at the point of contact. For each cycle, the force data were first adjusted to account for the finite sample thickness and an analytical solution for spherical indentation of a semi-infinite biphasic half space was fit to the adjusted force vs. displacement data^32^ via least squares minimization in MATLAB (version 9.5, The MathWorks, Inc., Natick, MA) with the contact modulus *E*_*c*_ and an offset displacement as free parameters. Unlike *F*_*pk*_, which describes the structural response over the entire test, *E*_*c*_ provides an estimate of the effective material behavior after the point of contact. Like *F*_*pk*_, *E*_*c*_ was normalized to *E*_*c*_ at cycle 3, the last cycle tested in the baseline 1X PBS bath (*E*_*c_PBS*_):

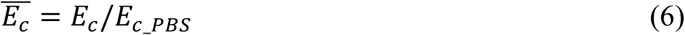

### Statistical analysis

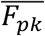 for the final exposure or recovery cycle and equilibrium time constants were analyzed using one-way analysis of variance (ANOVA) followed by Bonferroni’s post-hoc test for pairwise comparisons. Differences in *ε*_*osm*_ and 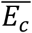 between conditions were analyzed using repeated measures ANOVA with Bonferroni’s post-hoc test for pairwise comparisons, as were differences in 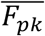 for the extended recovery meniscus experiment. Differences in t_50%_ between cartilage and meniscus for the 100% CA condition were assessed using an unpaired, two-tailed Student’s t-test. Analyses were performed using GraphPad Prism (version 8.0.2, GraphPad Software, La Jolla, CA) with significance set at p<0.05. Results are presented as mean ± 95% CI.

## Results

### Protocol validation

Cartilage (Fig. 2D) and meniscus (Fig. 2F) exhibited increased 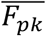 following exposure to 0.1X PBS and decreased 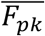 following exposure to 10X PBS, reflecting the known swelling and deswelling behaviors of these tissues. At the last exposure cycle (Fig. 2E; Fig. 2G), 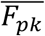 for the 0.1X PBS group was significantly greater than controls for both tissues while 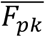 for the 10X PBS group was significantly lower than controls for cartilage. These results demonstrate that this testing protocol detects transient mechanical behaviors of cartilage and meniscus explants consistent with previous reports.

### Visual assessment of samples

Exposure to the contrast solutions, particularly when undiluted, altered the appearance of both cartilage (Fig. 3A) and meniscus (Fig. 3F) samples. While cartilage sample dimensions did not obviously change, meniscus samples in all contrast solutions distorted to some extent. Both tissues exhibited optical clearing (reduced opacity) with increased contrast agent concentration, evidence of fluid substitution consistent with the use of Omnipaque for tissue clearing in microscopy^33–35^.

**Figure 3:**
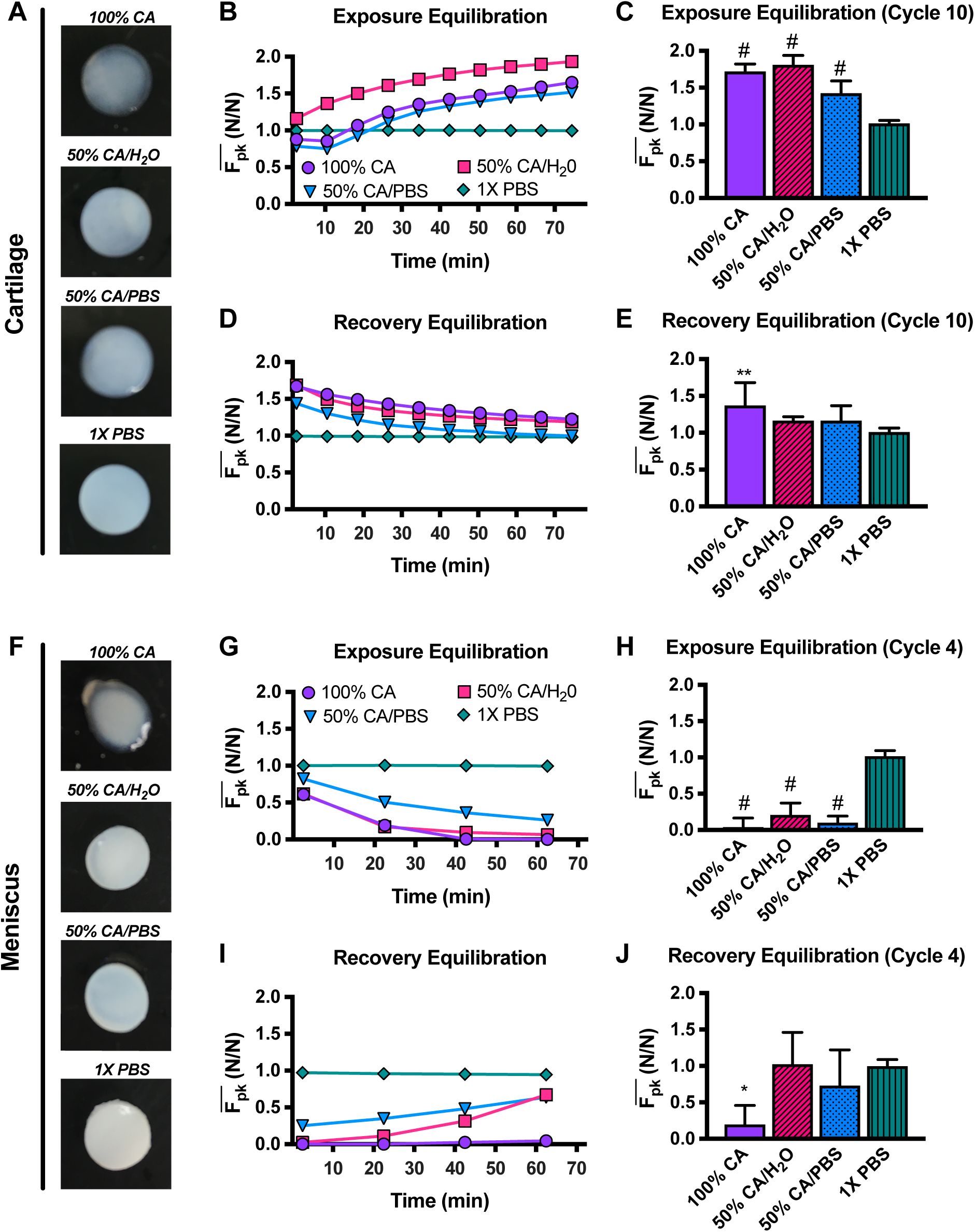
Exposure to contrast agents affects cartilage and meniscus appearance and differentially alters their response to indentation loading. Cartilage (A) and meniscus (F) explants equilibrated for 1h in 100% CA, or in a dilution of the contrast agent in water or PBS, lose their opacity when compared to the control explants equilibrated in PBS. Representative plots show that for cartilage explants, exposure to the different contrast agent dilutions resulted in long-term increase in the normalized peak force after 10 indentation cycles, with the 100% CA and 50% CA/PBS groups experiencing an initial decrease in normalized peak force in the first 2 cycles (B). Normalized peak force values on the last cycle (cycle 10) of indentation testing during exposure equilibration were all significantly greater than the PBS control (C). Re-equilibration in PBS resulted in an overall deswelling trend (D), with most groups returning to a baseline normalized peak force value except for the 100% CA group, which remained significantly greater than the control after 10 cycles of recovery (E). The opposite effects were observed in meniscus. During exposure equilibration (G), samples from all contrast agent groups experienced a rapid decrease in normalized peak force, with all groups maintaining significantly lower normalized peak force values than the PBS control on the last indentation cycle (H). Re-equilibration in PBS resulted in an increase in normalized peak force in meniscus (I), with most groups returning to baseline levels except for the 100% CA group which remained significantly lower than the control on the last cycle of recovery (J). n=4-5/group, * P<0.05, ** P<0.01, ± P<0.001, # P<0.0001.

### Normalized peak force values

Two cartilage and one meniscus sample were excluded from analysis due to inconsistencies in test results indicating procedural errors, with each experimental group maintaining at least four samples per condition. Patterns of 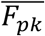 versus time indicate that the contrast solutions affected the two tissues differently, with distinctive patterns evident both among solutions and between tissues (Fig. 3B, 3D, 3G, 3I; Table 1A).

**Table 1:** Data summary. All results presented are mean ± 95% CI. (A) Normalized peak force values of cycle 10 of cartilage exposure and recovery equilibriums and cycle 4 of meniscus exposure and recovery equilibriums. Peak force values for exposure and recovery phases were normalized to the peak force value on the last indentation cycle during baseline testing. P-values for normalized peak force show significance in comparison to control groups (1X PBS). (B) Short-term (τ_short_), long-term (τ_long_), and recovery (τ_R_, τ_R_EXT_) equilibrium time constants for cartilage and meniscus samples obtained from exponential data fits. Short-term equilibrium time constants are only reported for 100% CA and 50% CA/PBS groups, which are the groups that exhibited the initial decrease in normalized peak force. Recovery equilibrium time constants for meniscus are only reported for extended tests. t_50%_ was determined as the time needed to reach 50% recovery in cartilage and meniscus samples initially treated with 100% CA. P-values for equilibrium time constants show significance in comparison to 100% CA groups, P-value for t_50%_ in meniscus shows significance in comparison to cartilage. (C) Normalized contact modulus for cartilage and meniscus samples. Contact modulus for exposure and recovery phases were normalized to the contact modulus on the last indentation cycle during baseline testing. P-values for normalized contact modulus show significance in comparison to control groups (1X PBS).

| A | Normalized Peak Force $\overline{F_{pk}}$ (N/N) | | | | | | | |
| --- | --- | --- | --- | --- | --- | --- | --- | --- |
|  | Cartilage |  |  |  | Meniscus |  |  |  |
|  | 100%<br>CA | 50%<br>CA/H<sub>2</sub>O | 50%<br>CA/PBS | PBS | 100%<br>CA | 50%<br>CA/H<sub>2</sub>O | 50%<br>CA/PBS | PBS |
| Exposure | 1.72±0.10 | 1.81±0.13 | 1.42±0.17 | 1.01±0.04 | 0.01±0.01 | 0.12±0.06 | 0.16±0.07 | 1.03±0.02 |
| P-value | <0.0001 | <0.0001 | <0.001 | --- | <0.0001 | <0.0001 | <0.0001 | --- |
| Recovery | 1.37±0.31 | 1.17±0.05 | 1.16±0.21 | 1.01±0.05 | 0.06±0.02 | 0.76±0.09 | 0.89±0.25 | 1.01±0.03 |
| P-value | 0.0020 | 0.2923 | 0.3868 | --- | 0.0147 | >0.9999 | >0.9999 | --- |

| B | Equilibrium Time Constants (s) |  |  |  |  |  |
| --- | --- | --- | --- | --- | --- | --- |
|  | Cartilage |  |  | Meniscus |  |  |
|  | 100%<br>CA | 50%<br>CA/H<sub>2</sub>O | 50%<br>CA/PBS | 100%<br>CA | 50%<br>CA/H<sub>2</sub>O | 50%<br>CA/PBS |
| $\tau_{\text{short}}$ | 321.7±249.2 | --- | 924.9±1539.9 | --- | --- | --- |
| P-value | --- | --- | >0.999 | --- | --- | --- |
| $\tau_{\text{long}}$ | 2343.0±1310.7 | 2443.8±775.6 | 2053.0±1276.9 | 1087.0±589.4 | 745.7±611.7 | 970.9±1039.7 |
| P-value | --- | >0.999 | >0.999 | --- | >0.999 | >0.999 |
| $\tau_{\text{R}}$ | 3971.9±2934.2 | 1457.8±286.3 | 2001.1±1554.8 | --- | --- | --- |
| P-value | --- | 0.0063 | 0.0327 | --- | --- | --- |
| $\tau_{\text{R\_EXT}}$ | --- | --- | --- | 4117.0±5078.7 | --- | --- |
| P-value | --- | --- | --- | --- | --- | --- |
| $t_{50\%}$ | 3672.0±2215.4 | --- | --- | 10155.3±5663.9 | --- | --- |
| P-value | --- | --- | --- | 0.01 | --- | --- |

| C | Normalized Contact Modulus $\overline{E_c}$ (MPa/MPa) | | | | | | | |
| --- | --- | --- | --- | --- | --- | --- | --- | --- |
|  | Cartilage |  |  |  | Meniscus |  |  |  |
|  | 100%<br>CA | 50%<br>CA/H<sub>2</sub>O | 50%<br>CA/PBS | PBS | 100%<br>CA | 50%<br>CA/H<sub>2</sub>O | 50%<br>CA/PBS | PBS |
| Exposure (Cycle 2) | 2.10±0.29 | 1.32±0.07 | 1.54±0.31 | 1.00±0.06 | --- | --- | --- | --- |
| P-value | 0.0023 | <0.0001 | 0.0309 | --- | --- | --- | --- | --- |
| Exposure (Last) | 2.47±0.89 | 1.75±0.18 | 1.89±0.45 | 1.02±0.05 | 0.11±0.18 | 0.24±0.32 | 0.16±0.09 | 1.00±0.11 |
| P-value | 0.0418 | 0.0005 | 0.0236 | --- | <0.0001 | 0.0012 | 0.0002 | --- |
| Recovery (Last) | 1.43±0.43 | 1.18±0.09 | 1.15±0.14 | 1.01±0.08 | 0.25±0.35 | 1.00±0.63 | 0.73±0.59 | 0.98±0.16 |
| P-value | 0.1577 | 0.0151 | 0.1356 | --- | 0.0010 | >0.9999 | 0.5494 | --- |

For cartilage, the hyperosmolal 100% CA and 50% CA/PBS groups showed a transient, short-term decrease in 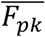 not seen in the roughly isosmolal 50% CA/H_2_O group, but all contrast groups exhibited long-term increases in 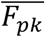 that appeared to approach equilibrium at the end of the exposure phase (Fig. 3B). 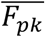 on the last exposure cycle (cycle 10) was significantly greater for all contrast agent groups than for the PBS controls (Fig. 3C). Upon returning the baths to 1X PBS, 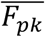 for cartilage contrast groups decayed towards their baseline levels (Fig. 3D), with all samples approaching equilibrium and only the 100% CA group remaining significantly greater than controls at the last recovery cycle (Fig. 3E).

The effects on meniscal tissue were strikingly different. All contrast solutions induced substantial decreases in meniscus 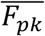 following exposure (Fig. 3G). 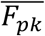 for all meniscus contrast groups were significantly lower than controls at the last exposure cycle (Fig. 3H). 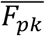 considerably increased for the diluted contrast groups during recovery (Fig. 3I) but remained close to zero for the 100% CA group, significantly lower than controls at the last recovery cycle (Fig. 3J).

To further assess recovery of meniscus samples exposed to 100% CA, additional samples were monitored for 4 exposure cycles plus an extended period of 25 recovery cycles to provide an extra 6 hours of re-equilibration in 1X PBS. Meniscal samples reached baseline 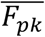 after approximately 10 cycles but 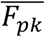 continued to increase with time (Fig. 4). Interestingly, 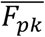 after 15 cycles of recovery was significantly greater for meniscal samples previously treated with 100% CA than for controls, suggesting that structural effects resulting from exposure to contrast agents on meniscus linger for hours after removal of the contrast agent.

**Figure 4:**
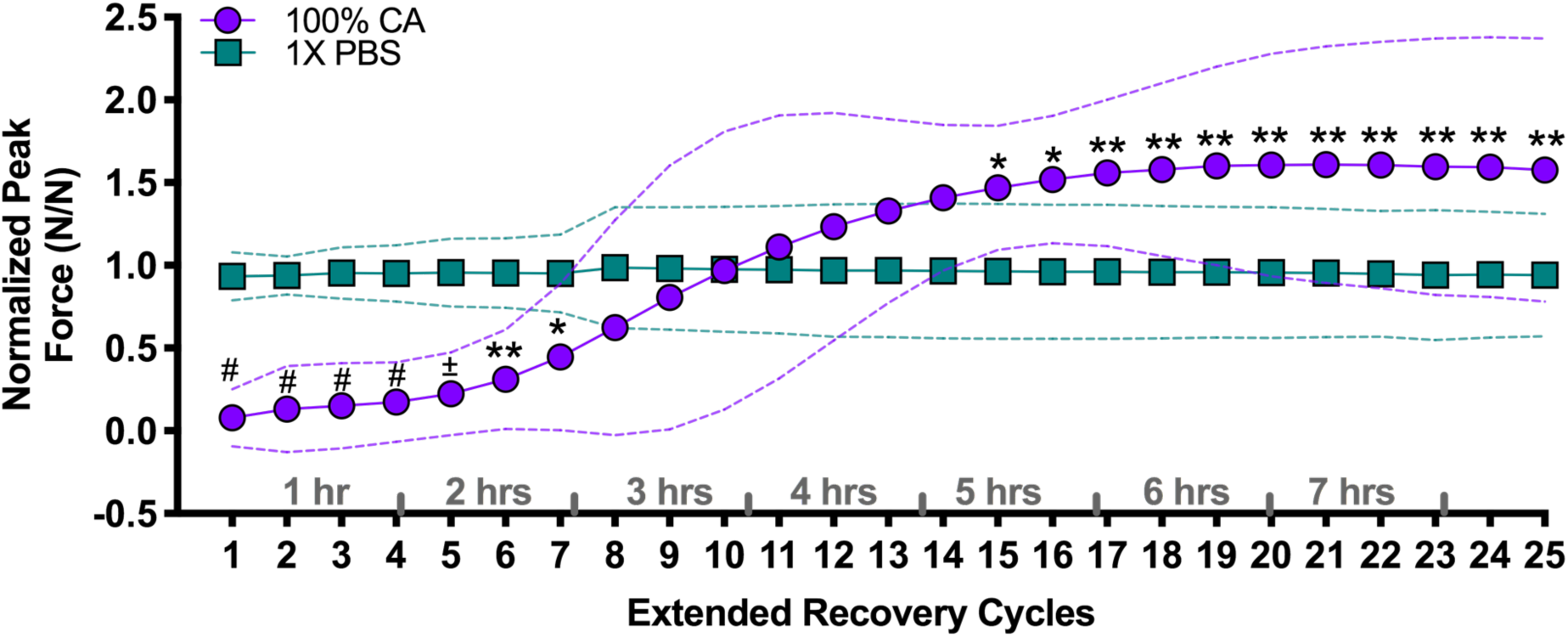
Meniscus normalized peak force considerably increases with extended re-equilibration times. A subset of meniscus explants was subjected to an extended recovery indentation test consisting of 25 cycles. Meniscus explants reached baseline normalized peak force values after 8 cycles (∼2.5 hours) in PBS and continued increasing way beyond this baseline level. Normalized peak force values after cycles 1-7 of recovery were significantly lower than the PBS control, while values after cycle 15 of recovery were significantly larger than the PBS control. n=3/group, * P<0.05, ** P<0.01, ± P<0.001, # P<0.0001.

### Equilibration time constants

Most testing conditions for cartilage and meniscus samples resulted in behaviors well described by exponential fits, with no significant differences in *τ*_*E*_ among contrast solutions for either tissue (Fig. 5A). The additional equilibration time constant *τ*_*short*_, describing the initial decrease in 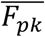 for cartilage samples equilibrated in 100% CA or 50% CA/PBS, did not significantly differ between groups (Fig. 5A). For cartilage, the recovery time constant for the 100% CA group was significantly greater than for the 50% CA/H_2_O and 50% CA/PBS groups (Fig. 5B). To compare the time response of recovery for cartilage and meniscus samples previously equilibrated in 100% CA, the times required to reach 50% recovery were tabulated. Meniscus required approximately 3-times longer to reach 50% recovery than cartilage (Fig. 5C). Table 1B summarizes all equilibrium time constant results. Individual exponential fits for each cartilage and meniscus sample are included as part of the supplemental data (Cartilage: Fig. S3; Meniscus: Fig. S4).

**Figure 5:**
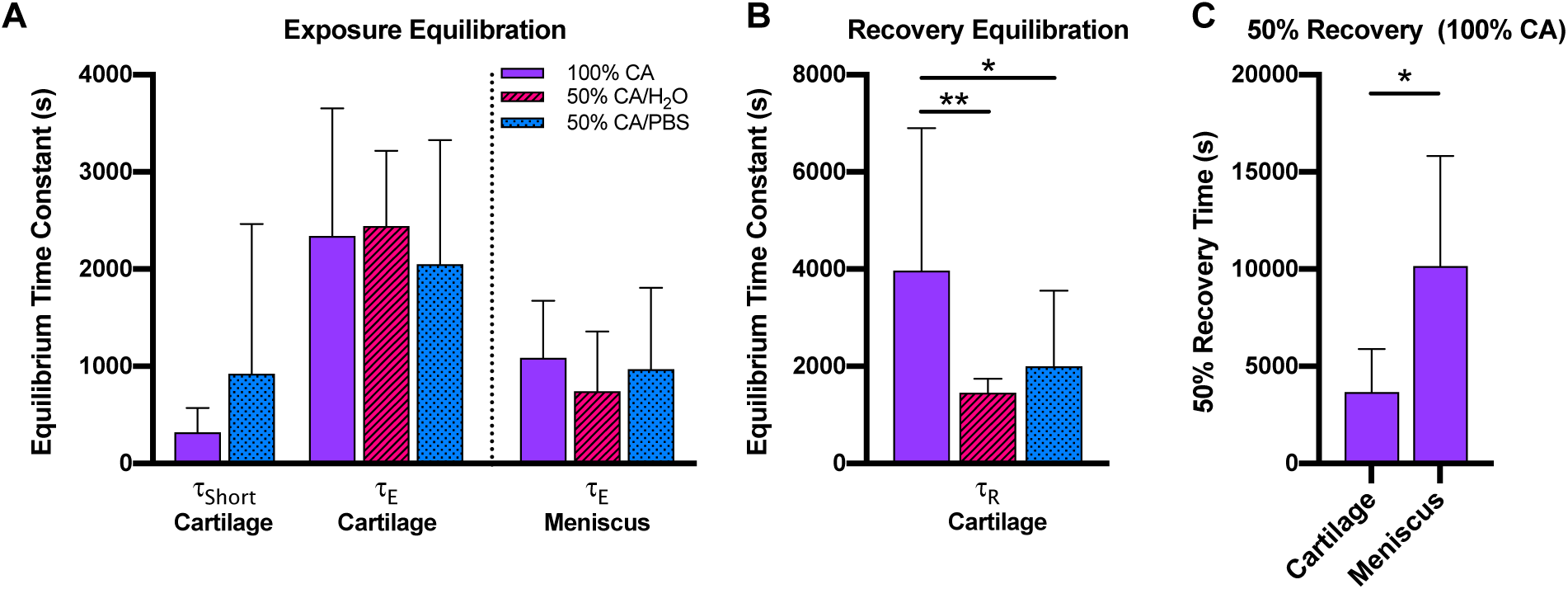
Meniscus requires longer recovery times to return to baseline equilibrium. (A) No significant differences were detected in equilibration time constant *τ*_*short*_ (representing the initial decrease in normalized peak force) between cartilage groups. No significant differences in equilibration time constant *τ*_*E*_ (representing the long-term normalized peak force behavior) were observed between the CA groups for both cartilage and meniscus samples. (B) The recovery equilibration time constant for cartilage, τ_R_, was significantly longer for the 100% CA group than both 50% CA/H_2_O and 50% CA/PBS groups. (C) Time required to reach 50% recovery after exposure to 100% CA was significantly greater for meniscus than cartilage. Results are mean + 95% CI, n=3-5/group, *P<0.05, **P<0.01.

### Changes to contact stiffness

In cartilage samples exposed to 100% CA or 50% CA/PBS, *ε*_*osm*_ trends exhibited an initial decrease followed by a long-term increase with values that remained negative for the duration of the test (Fig. 6A), suggesting the effect of the initial tissue contraction (negative *ε*_*osm*_) persisted as samples experienced subsequent relative expansion. During recovery equilibration, samples assigned to 100% CA and 50% CA/PBS groups continued expanding, with *ε*_*osm*_ eventually equilibrating close to baseline levels. Cartilage equilibrated in 50% CA/H_2_O showed minimal changes in *ε*_*osm*_, with no significant differences detected from the control group. 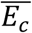 in cartilage (Fig. 6B) consistently increased for all contrast agent groups during exposure equilibration, suggesting a stiffening effect. 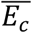 decreased during recovery equilibration for all groups, with only the 50% CA/H_2_O group significantly larger than the PBS control at the end of recovery testing.

**Figure 6:**
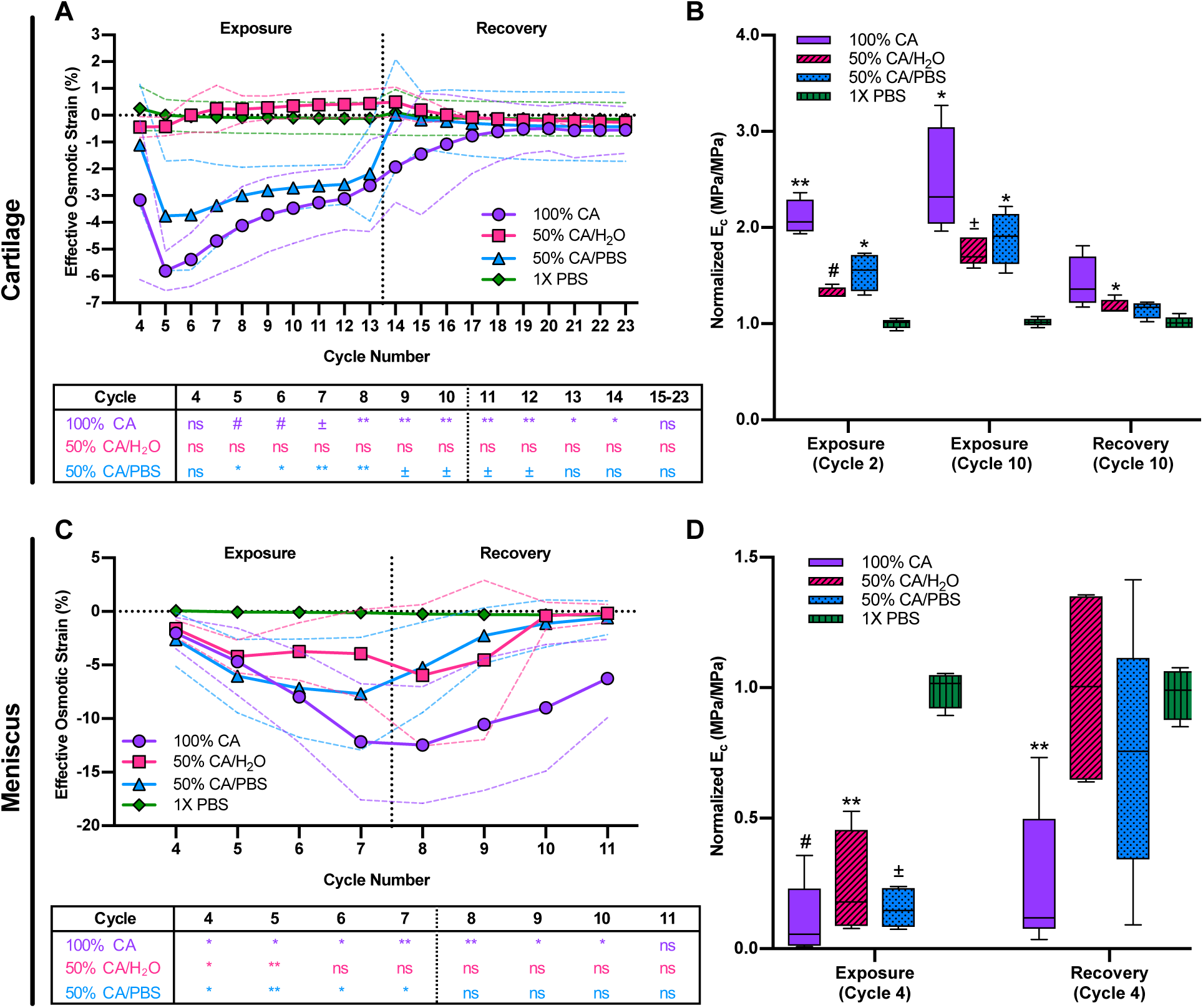
Contrast agent dilutions stiffen cartilage and soften meniscus at the surface. Changes in the effective osmotic strain *ε*_*osm*_ of (A) cartilage samples showed that exposure to 100% CA and 50% CA/PBS resulted in an initial softening or contraction effect at the surface with the most negative value of *ε*_*osm*_ at cycle 2 of exposure equilibration. After cycle 2, samples exhibited an increasing trend in osmotic strain. The 50% CA/H_2_O group exhibited an increasing, positive *ε*_*osm*_, indicating swelling/expansion of the samples. Interestingly, the significant increases in normalized effective contact modulus E_c_ of cartilage samples (B) suggest that the long-term trends resulting from changes in ionic strength, rather than osmolality, govern the tissue response at the bulk level. In meniscus, exposure equilibration lead to a rapid decrease in the effective osmotic strain (C), consistent with the normalized peak force data (Fig. 3G) and suggesting a deswelling effect in meniscus samples. The significant decreases in normalized effective contact modulus for meniscus (D) also indicate a softening effect, which remains significant even after 4 cycles of recovery for the 100% CA group. Dashed lines in effective osmotic strain plots represent 95% CI. In normalized effective contact modulus plots, upper and lower box boundaries represent 25^th^ and 75^th^ percentiles with the middle line representing the mean, whiskers show 5^th^ and 95^th^ percentile. n=4-5/group, significance set with respect to PBS control groups *P<0.05, **P<0.01, ±P<0.001, #P<0.0001.

All meniscus contrast agent groups exhibited decreasing *ε*_*osm*_ trends that were reversed during recovery equilibration (Fig. 6C). Consistent with *ε*_*osm*_, 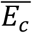 was lower for all contrast agent groups in meniscus compared to control and returned to baseline levels after 4 cycles of recovery, except for the 100% CA group which remained significantly lower than the control (Fig. 6D). These results suggest reduced stiffness with exposure to contrast agents in meniscus.

## Discussion

CT arthrography protocols vary widely, with parameters such as contrast agent dilution being based primarily on their effects on image quality (i.e. iodine concentration sufficient to ensure image clarity 30-60 minutes after injection, but low enough to avoid major beam hardening artifacts). This study characterized the transient response of cartilage and meniscus tissue samples exposed to various dilutions of iohexol, a non-ionic, iodinated, low-osmolar contrast agent commonly used for CT arthrography. We found that clinically relevant contrast agent formulations did, in fact, alter the mechanical response of joint tissues in a time frame relevant to clinical scan protocols. Surprisingly, cartilage and meniscus tissue samples exhibited opposing behaviors when exposed to equivalent dilutions of the contrast agent. Our findings suggest that exposure to non-ionic, iodinated contrast agents could render cartilage and meniscus susceptible to load-induced material or biological damage during arthrography procedures, and that effects could linger for hours after contrast administration. Note that reductions of contrast agent concentration due to physiologic clearance are not step changes, and more gradual reductions would be expected to prolong the after-effects of contrast exposure.

Cartilage explants exhibited an initial decrease in 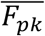 when equilibrated in 100% CA and 50% CA/PBS groups, as well as a long-term increase in 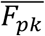 that was present in every contrast agent group. We hypothesize that these trends reflect two competing phenomena that occur simultaneously. The hyperosmotic nature of these contrast solutions induces an immediate deswelling effect, as the reduced chemical potential of water in the bath draws fluid out of the tissue. However, the low ion content of the contrast agent solutions (particularly the 100% CA and 50% CA/H_2_O groups) substantially perturbs the electrochemical interactions among the negatively-charged cartilage matrix, the interstitial ions, and the ions in the bath, causing an efflux of ions from the tissue and resulting in an increased osmotic swelling stress^36^. As the non-ionic contrast molecule (the primary neutral solute in these mixtures) is able to enter the tissue and would eventually equilibrate, the long-term response is expected to reflect the low ionic strength of these hyperosmolal solutions. In the 100% CA and 50% CA/PBS cartilage groups, the high osmolality initially dominates the mechanical response, but the long-term response of all cartilage contrast agent groups are dominated by the low ionic strength (Fig. 7A).

**Figure 7:**
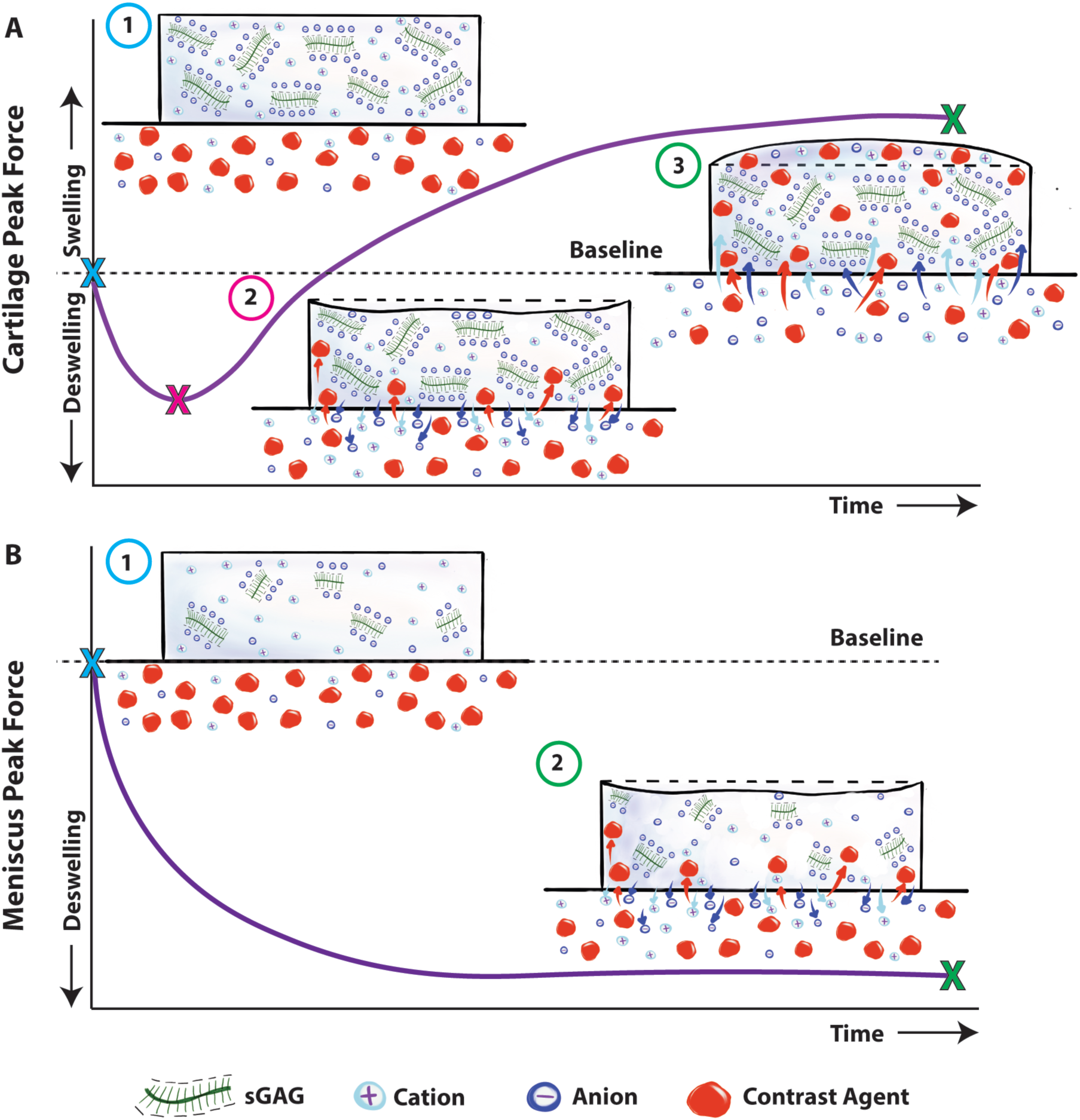
Schematic representation of competing hyperosmotic and hypotonic effects in cartilage and meniscus samples. A) Starting at an equilibrium state in 1X PBS (1), cartilage samples exposed to 100% CA experience an initial deswelling behavior (2) that is governed by the hyperosmolality of the contrast agent bath, with the high solute (contrast agent) concentration in the bath solution drawing fluid out of the samples. As contrast agent molecules diffuse into the tissue, this effect is reduced and the lack of ions in the contrast agent bath (i.e. the hypotonicity of the bath) dominates the subsequent response, drawing ions out of the tissue and driving fluid into the tissue as it moves towards a new equilibrium (3). B) The initial response of meniscus samples is similarly governed by the hyperosmolality of the contrast agent bath, drawing fluid out of the tissue sample. Due to the relatively low fixed charge density of meniscal tissue, however, electrochemical interactions between the tissue matrix and the fluid are much weaker, and the effects of a hypotonic bath solution are relatively small. As a result, the long-term equilibrium response is consistent with the initial deswelling induced by the hypoosmolal bath.

While the same phenomena affect meniscus tissue, the much lower sGAG concentration^37^ compared to cartilage substantially reduced the impact of the solution ionic strength. Consequently, the responses of meniscus explants appeared to be dominated by the deswelling behavior characteristic of exposure to a hyperosmolal solution (Fig. 7B). The iohexol formulation used with meniscus samples was less hyperosmolal than that used with cartilage (Fig. 2H), and we would expect an even larger discrepancy between tissues if identical formulations were used. Curiously, substantial deswelling was observed even in the slightly hypoosmolal 50%CA/H_2_O group for meniscus, suggesting either more complex interactions (e.g. involving intrafibrillar water within collagen fibrils) or unknown microstructural changes induced by the contrast molecules.

For cartilage, these trends were largely transient, with deswelling and a return to baseline behaviors in 50% CA/PBS and 50% CA/H_2_O groups within an hour of recovery in PBS. The 100% CA group also returned towards baseline behavior but remained significantly swollen after an hour of recovery. For meniscus, the 50% CA/PBS and 50% CA/H_2_O groups also exhibited substantial recovery (reswelling) but did not fully recover within an hour. The 100% CA group for meniscus exhibited minimal initial recovery and required 3 hours to return to baseline, with an eventual and sustained increase in 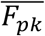 to levels above baseline. This may indicate an unexpected effect of the contrast agent on the meniscal microstructure.

While an important first step in determining the mechanical effects of contrast agents in cartilage and meniscus surfaces, this study has some limitations. In vivo, joints experience a more complex and highly variable range of physiological loading that is difficult to replicate ex-vivo. Harvesting of samples, particularly in the case of meniscus tissues, disrupts the collagen network and potentially alters the mechanical behavior. Nonetheless, preliminary tests on intact meniscus specimens performed while developing the indentation testing protocol revealed minimal effects of explant extraction on the mechanical response captured by these indentation tests. Juvenile bovine tissues are less mature than their adult human counterparts and may differ in relevant aspects of composition including collagen and GAG content, collagen cross-linking, etc. that would be expected to affect the relative degree of mechanical alterations due to contrast exposure. The persistent alterations observed in meniscal behavior, including the increased 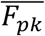 during long-term recovery, allude to potential structural changes that should be addressed in future studies on human tissues. It is important to note that clinical arthrography is not typically performed in healthy joints, and the effects of contrast agents on diseased human tissues, including ones with compromised surface layers that allow more rapid transport at the tissue-fluid interface, will be important to evaluate going forward. Although our study did not include full-thickness explants, the use of partial thickness samples with intact surface layers is appropriate for studies of relative short-term, transient responses such as this one. Nevertheless, further studies examining tissues in the context of intact joints (cadaveric or living) will provide more directly relevant insights. Preliminary vital dye staining of cartilage showed minimal cytotoxicity after 1hr exposure to 100% CA (Fig. S5); however, the large step change in the osmotic environment associated with contrast injection suggests that more thorough evaluations of biological effects, including biochemical and histological analysis, would be prudent.

The work presented here reveals that exposure to non-ionic, iodinated contrast agents at clinically relevant dilutions can rapidly and substantially alter the mechanical behaviors of cartilage and meniscus. Given the strong nature of the ionic interactions that govern the swelling behaviors of cartilage and meniscus, we posit that the effects of contrast agent media with larger osmolality differences (compared to physiological conditions) will be even stronger on the mechanical response of these tissues, but that producing that mismatch with ionic and nonionic agents will result in different tissue responses. Although experimental studies would be required for verification, we surmise that exposure to iso-osmotic, non-ionic contrast agents such as iodixanol would still produce a swelling response in cartilage due to low ionic strength but would not produce the initial deswelling related to the high osmolality of iohexol, while the meniscus response might be less substantial than reported here. Ionic contrast solutions are predominantly both hyperosmolal and hypotonic and undiluted ionic contrast solutions would thus be expected to induce deswelling and softening in both cartilage (as previously observed with Hexabrix^29^) and meniscus, rendering the surfaces of both tissues more susceptible to physical and biological damage upon loading.

Given the rapid changes in tissue behaviors observed in these studies, it may be prudent to limit common clinical practices such as pre-scan patient ambulation and instead favor gentler passive joint movement^38^ as a way to distribute contrast while minimizing potential tissue damage. In a similar manner, the long recovery time (particularly for the fibrocartilaginous meniscus) upon complete removal of the contrast agent raises concerns about the timing of patient re-ambulation following the procedure. Our results advise the establishment of new guidelines for CT arthrography that consider not only systemic reactions and image quality but also the unique physical effects of contrast agents on hydrated joint tissues.

## Acknowledgements

The authors thank Dr. Alberto Arvayo for assistance in preparing Fig. 7.

## Author contributions

Dr. Baylon contributed to the concept and design of the study, data acquisition, analysis and interpretation of results and drafting of the manuscript. Ms. Crowder assisted with data collection. Drs. Gold and Levenston contributed to the conception and design of the study, analysis and interpretation of results, and provided funding for the study. All authors critically revised the article and approved the final version submitted. Dr. Baylon and Dr. Levenston take responsibility for the integrity of the work presented and can be contacted at, respectively.

## References

1. Peterson JJ, Bancroft LW. History of Arthrography. Radiol Clin North Am. 2009;47(3):373–386. doi: 10.1016/j.rcl.2008.12.001

2. Carter K, Mudigonda S. Arthrography and Injection Procedures. Imaging Arthritis Metab Bone Dis. 2009;5:60–80. doi: 10.1016/B978-0-323-04177-5.00005-7

3. Hirvasniemi J, Kulmala KAM, Lammentausta E, Ojala R, Lehenkari P, Kamel A, et al. In vivo comparison of delayed gadolinium-enhanced MRI of cartilage and delayed quantitative CT arthrography in imaging of articular cartilage. Osteoarthritis Cartilage. 2013;21(3):434–442. doi: 10.1016/j.joca.2012.12.009

4. van Tiel J, Siebelt M, Reijman M, Bos PK, Waarsing JH, Zuurmond AM, et al. Quantitative in vivo CT arthrography of the human osteoarthritic knee to estimate cartilage sulphated glycosaminoglycan content: correlation with ex-vivo reference standards. Osteoarthr Cartil. 2016;24(6):1012–1020. doi: 10.1016/j.joca.2016.01.137

5. West A, Marshall T, Bearcroft P. CT of the musculoskeletal system: What is left is the days of MRI? Eur Radiol. 2009;19:152–164. doi: 10.1007/s00330-008-1129-0

6. Kalke R, Di Primio G, Schweitzer M. MR and CT Arthrography of the Knee. Semin Musculoskelet Radiol. 2012;16(01):057–068. doi: 10.1055/s-0032-1304301

7. Lusic H, Grinstaff MW. X-ray-computed tomography contrast agents. Chem Rev. 2013;113(3):1641–1666. doi: 10.1021/cr200358s

8. Mcclennan BL. Preston M. Hickey memorial lecture. Ionic and Nonionic lodinated Memorial Lecture Contrast Media: Evolution and Strategies for Use. Am J Roentgenol. 1990;155(2):225–233. http://www.ajronline.org. Accessed September 13, 2018.

9. Katayama H, Yamaguchi K, Kozuka T, Takashima T, Seez P, Matsuura K. Adverse reactions to ionic and nonionic contrast media. A report from the Japanese Committee on the Safety of Contrast Media. Radiology. 1990;175(3):621–628. doi: 10.1148/radiology.175.3.2343107

10. Baumgarten M, Bloebaum RD, Ross SD, Campbell P, Sarmiento A. Normal human synovial fluid: osmolality and exercise-induced changes. J Bone Joint Surg Am. 1985;67(9):1336–1339. http://www.ncbi.nlm.nih.gov/pubmed/4077904. Accessed January 19, 2019.

11. Newman PJ, Grana WA. The changes in human synovial fluid osmolality associated with traumatic or mechanical abnormalities of the knee. Arthrosc J Arthrosc Relat Surg. 1988;4(3):179–181. doi: 10.1016/S0749-8063(88)80023-9

12. Shanfield S, Campbell P, Baumgarten M, Bloebaum R, Sarmiento A. Synovial fluid osmolality in osteoarthritis and rheumatoid arthritis. Clin Orthop Relat Res. 1988;(235):289–295. http://www.ncbi.nlm.nih.gov/pubmed/3416536. Accessed March 14, 2019.

13. Omoumi P, Mercier GA, Lecouvet F, Simoni P, Vande Berg BC. CT Arthrography, MR Arthrography, PET, and Scintigraphy in Osteoarthritis. Radiolodic Clin North Am. 2009;47:595–615. doi: 10.1016/j.rcl.2009.04.005

14. Rastogi AK, Davis KW, Ross A, Rosas HG. Fundamentals of Joint Injection. Am J Roentgenol. 2016;207(3):484–494. doi: 10.2214/AJR.16.16243

15. Department of Radiology University Of Wisconsin School of Medicine and Public Health. Muskuloskeletal Imaging and Intervention Section | Imaging Procedures Knee MR or CT Arthogram. https://www.radiology.wisc.edu/sections/msk/interventional/Knee_CT_arthrogram/index. php. Published 2018. Accessed March 2, 2018.

16. Choi J-H, Mcwalter EJ, Datta S, Mueller K, Maier A, Gold GE, et al. In Vivo 3D Measurement of Time-dependent Human Knee Joint Compression and Cartilage Strain During Static Weight-Bearing. In: Proceedings of the Orthopaedic Research Society (ORS) Annual Meeting. Orlando, FL; 2016. https://www.ors.org/Transactions/62/0120.pdf. Accessed March 2, 2018.

17. Holland P, Davies AM, Cassar-Pullicino V. Computed Tomographic Arthrography in the Assessment of Osteochondritis Dissecans of the Elbow. Clin Radiol. 1994;49:231–235. https://ac-els-cdn-com.laneproxy.stanford.edu/S000992600581846X/1-s2.0-S000992600581846X-main.pdf?_tid=6bf1bd13-25c5-4ab5-b38b-1a49339207b2&acdnat=1520015142_c6593d83dc5ae9ae67d4ca209b7ffd55. Accessed March 2, 2018.

18. Hao Y, Cui F, Zhu W, Lu L, Wang Y. Evaluation of the Clinical Significance of Classification of Traumatic Anterior Shoulder Instability using Double-contrast Computed Tomography Arthrography. J Int Med Res. 2011;39:424–434. http://journals.sagepub.com/doi/pdf/10.1177/147323001103900210. Accessed March 2, 2018.

19. Farsø Nielsen F, de Carvalho A, Hjøllund Madsen E. Omnipaque and urografin in arthrography of the knee. Acta Radiol Diagn (Stockh). 1984;25(2):151–154. http://www.ncbi.nlm.nih.gov/pubmed/6731021. Accessed October 24, 2018.

20. Kirschke JS, Braun S, Baum T, Holwein C, Schaeffeler C, Imhoff AB, et al. Diagnostic Value of CT Arthrography for Evaluation of Osteochondral Lesions at the Ankle. Biomed Res Int. 2016;2016:1–11. doi: 10.1155/2016/3594253

21. Eisenberg SR, Grodzinsky AJ. Swelling of Articular Cartilage and Other Connective Tissues: Electromechanochemical Forces. J Orthop Res. 1985;3(2):148–159. http://www.ncbi.nlm.nih.gov/pubmed/3998893.

22. Nguyen AM, Levenston ME. Comparison of Osmotic Swelling Influences on Meniscal Fibrocartilage and Articular Cartilage Tissue Mechanics in Compression and Shear. J Orthop Res. 2012;30(1):95–102. doi: 10.1002/jor.21493

23. Sun DD, Guo XE, Likhitpanichkul M, Lai WM, Mow VC. The Influence of the Fixed Negative Charges on Mechanical and Electrical Behaviors of Articular Cartilage Under Unconfined Compression. J Biomech Eng. 2004;126(1):6. doi: 10.1115/1.1644562

24. Korhonen RK, Jurvelin JS. Compressive and tensile properties of articular cartilage in axial loading are modulated differently by osmotic environment. Med Eng Phys. 2010;32:155–160. doi: 10.1016/j.medengphy.2009.11.004

25. Bursac P, Arnoczky S, York A. Dynamic compressive behavior of human meniscus correlates with its extra-cellular matrix composition. Biorheology. 2009;46(3):227–237. doi: 10.3233/BIR-2009-0537

26. Andrews SHJ, Rattner JB, Shrive NG, Ronsky JL. Swelling significantly affects the material properties of the menisci in compression. J Biomech. 2015;48(8):1485–1489. doi: 10.1016/j.jbiomech.2015.02.001

27. Wilson CG, Vanderploeg EJ, Zuo F, Sandy JD, Levenston ME. Aggrecanolysis and in vitro matrix degradation in the immature bovine meniscus: mechanisms and functional implications. Arthritis Res Ther. 2009;11(6):R173. doi: 10.1186/ar2862

28. Baylon EG, Levenston ME. Osmotic Swelling Responses are Conserved Across Cartilaginous Tissues with Varied Sulfated-Glycosaminoglycan Contents. J Orthop Res. November 2019. doi: 10.1002/jor.24521

29. Turunen MJ, Töyräs J, Lammi MJ, Jurvelin JS, Korhonen RK. Hyperosmolaric contrast agents in cartilage tomography may expose cartilage to overload-induced cell death. J Biomech. 2012;45(3):497–503. doi: 10.1016/J.JBIOMECH.2011.11.049

30. U.S. Food & Drug Administration. Drugs@FDA: FDA Approved Drug Products. https://www.accessdata.fda.gov/scripts/cder/daf/index.cfm?event=overview.process&ApplNo=018956. Published 2018. Accessed November 9, 2018.

31. Baylon EG, Levenston ME. Osmotic Swelling Responses are Conserved Across Cartilaginous Tissues with Varied Sulfated-Glycosaminoglycan Contents. bioRxiv. November 2018:459115. doi: 10.1101/459115

32. Moore AC, Zimmerman BK, Chen X, Lu XL, Burris DL. Experimental characterization of biphasic materials using rate-controlled Hertzian indentation. Tribol Int. 2015;89:2–8. doi: 10.1016/j.triboint.2015.02.001

33. Sdobnov AY, Darvin ME, Genina EA, Bashkatov AN, Lademann J, Tuchin VV. Recent progress in tissue optical clearing for spectroscopic application. Spectrochim Acta Part A Mol Biomol Spectrosc. 2018;197:216–229. doi: 10.1016/J.SAA.2018.01.085

34. Alexandrovskaya YM, Evtushenko EG, Obrezkova MM, Tuchin VV, Sobol EN. Control of optical transparency and infrared laser heating of costal cartilage via injection of iohexol. J Biophotonics. 2018;11(12):e201800195. doi: 10.1002/jbio.201800195

35. Ke M-T, Nakai Y, Fujimoto S, Takayama R, Yoshida S, Kitajima TS, et al. Super-Resolution Mapping of Neuronal Circuitry With an Index-Optimized Clearing Agent. Cell Rep. 2016;14(11):2718–2732. doi: 10.1016/J.CELREP.2016.02.057

36. Mattern KJ, Nakornchai C, Deen WM. Darcy Permeability of Agarose-Glycosaminoglycan Gels Analyzed Using Fiber-Mixture and Donnan Models. Biophys J. 2008;95(2):648–656. doi: 10.1529/biophysj.107.127316

37. Mow VC, Huiskes R. Basic Orthopaedic Biomechanics & Mechano-Biology. Philadelphia: Williams & Wilkins; 2005.

38. van Tiel J, Siebelt M, Waarsing JH, Piscaer TM, van Straten M, Booij R, et al. CT arthrography of the human knee to measure cartilage quality with low radiation dose. Osteoarthr Cartil. 2012;20(7):678–685. doi: 10.1016/J.JOCA.2012.03.007

