## Supplemental Data for "Non-ionic CT Contrast Solutions Rapidly Alter Bovine Cartilage and Meniscus Mechanics"

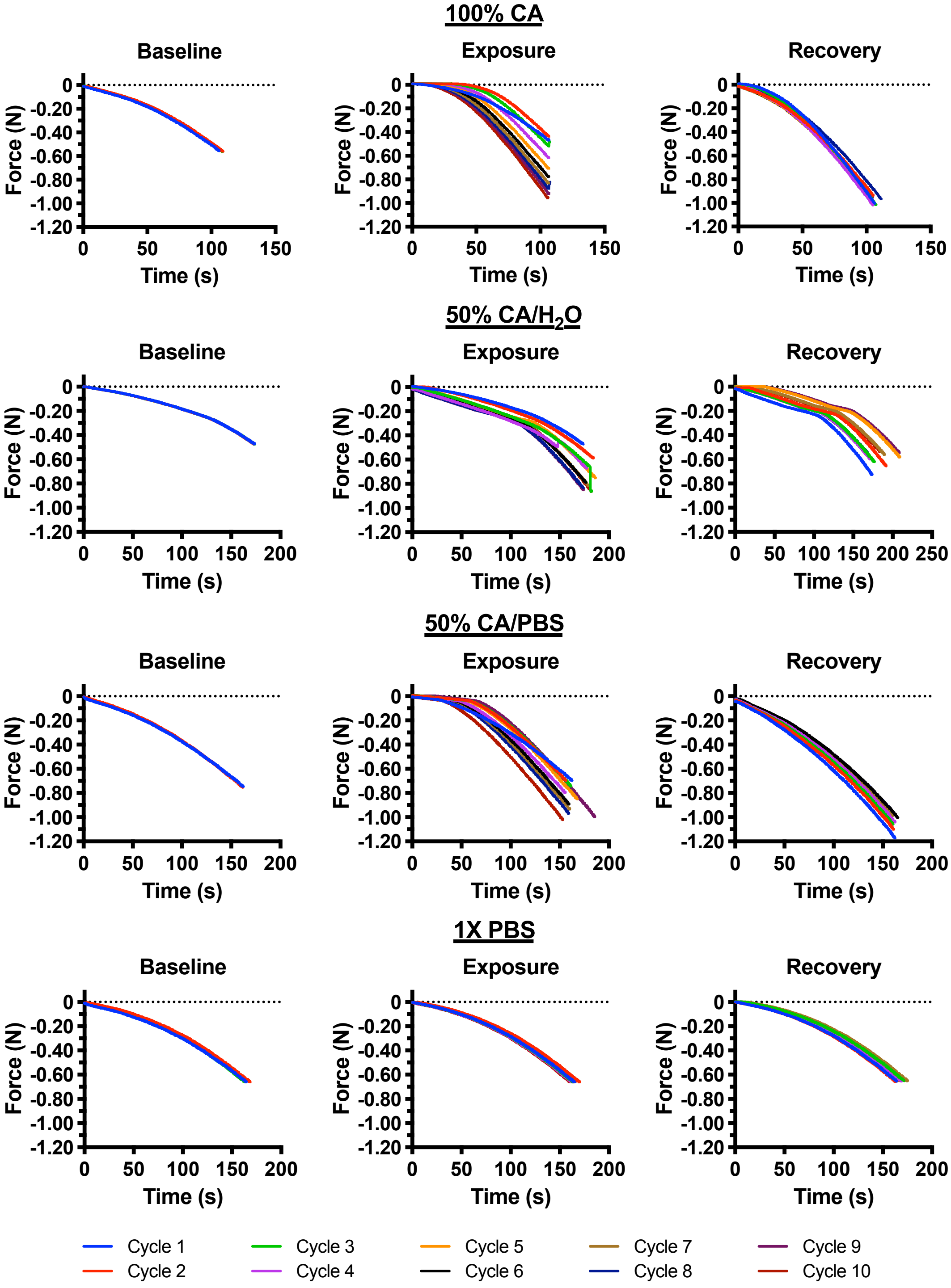


**Figure S1: Representative cartilage loading curves.** Qualitatively, these plots show that exposure to 1X PBS has a negligible effect in the cartilage response to loading. Equilibration in 100% CA and 50% CA/PBS results in an initial decrease in the slope of the loading curves – followed by a gradual increase, consistent with the results obtained from the contact model. Cartilage recovery curves show a smaller effect with the slope of the curves gradually decreasing as the surface softens to return to baseline.

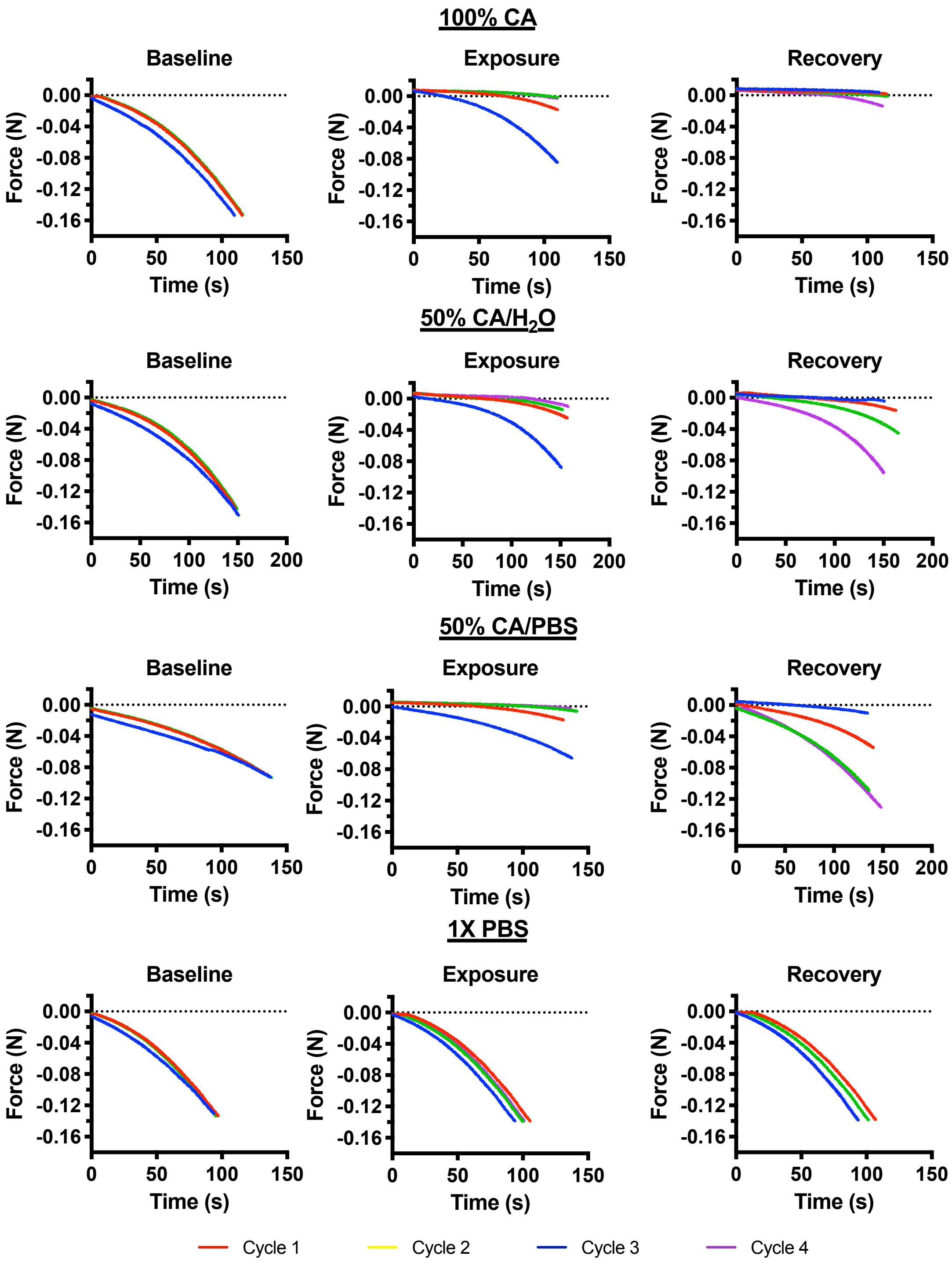


**Figure S2: Representative meniscus loading curves.** Qualitatively, these plots show that exposure to 1X PBS has a negligible effect in the meniscus response to loading. Contrast agent equilibration leads to a rapid decrease in the slope of the loading curves, particularly of the 100% CA and 50% CA/PBS groups, consistent with the results obtained from the contact model. During recovery equilibration, meniscus explants previously equilibrated in 100% CA fail to recover, while groups equilibrated in CA dilutions return to baseline 1X PBS values.

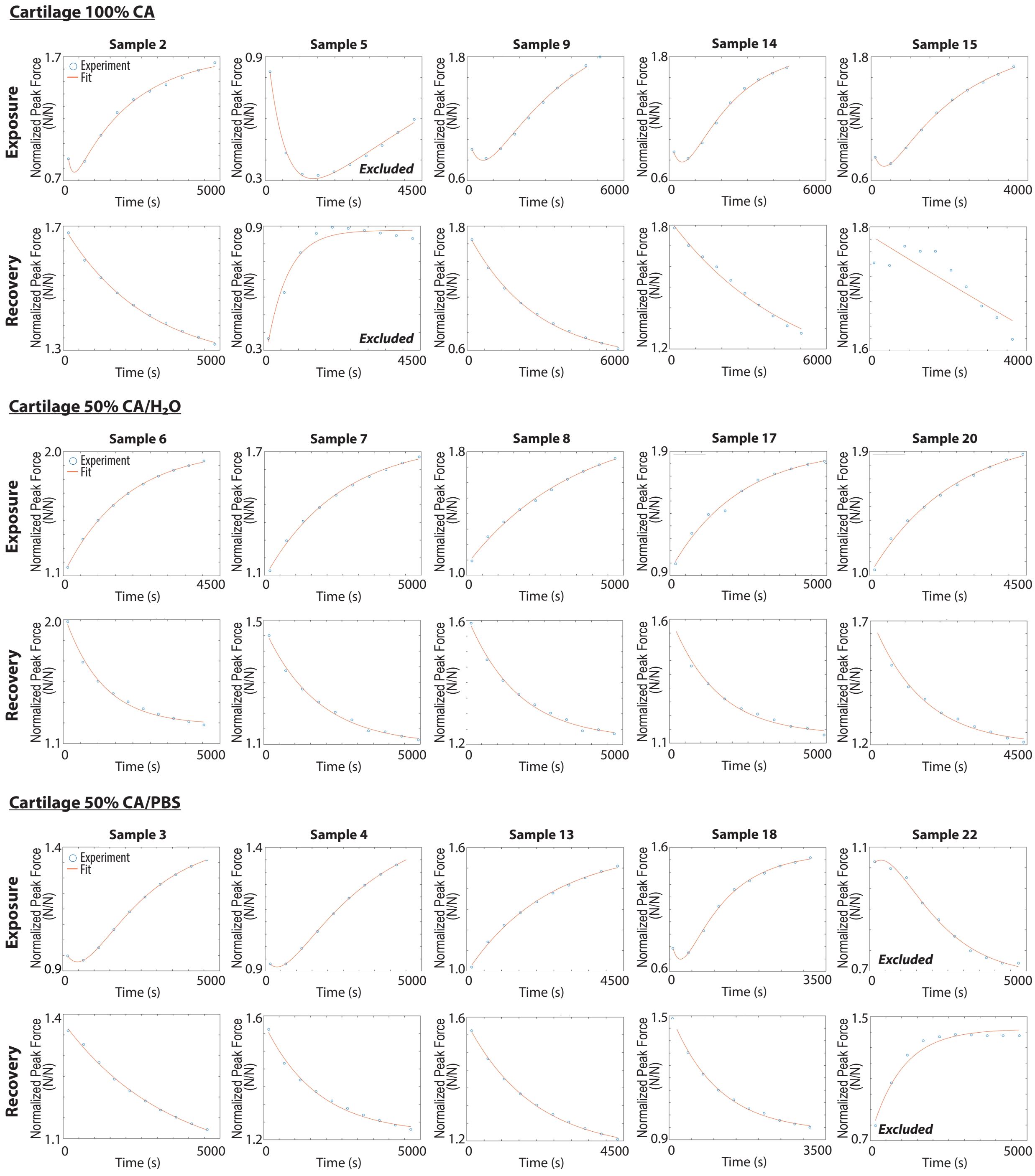


**Figure S3: Individual cartilage contrast agent and recovery data fits.** Blue markers represent raw data (normalized peak force), red line represents data fit. A biexponential data fit was used for 100% CA and 50% CA/PBS groups while a monoexponential fit was used for 50% CA/$H_2O$ and all recovery data. Samples 5 (100% CA) and 22 (50% CA/PBS) were excluded due to inconsistencies in their mechanical behavior.

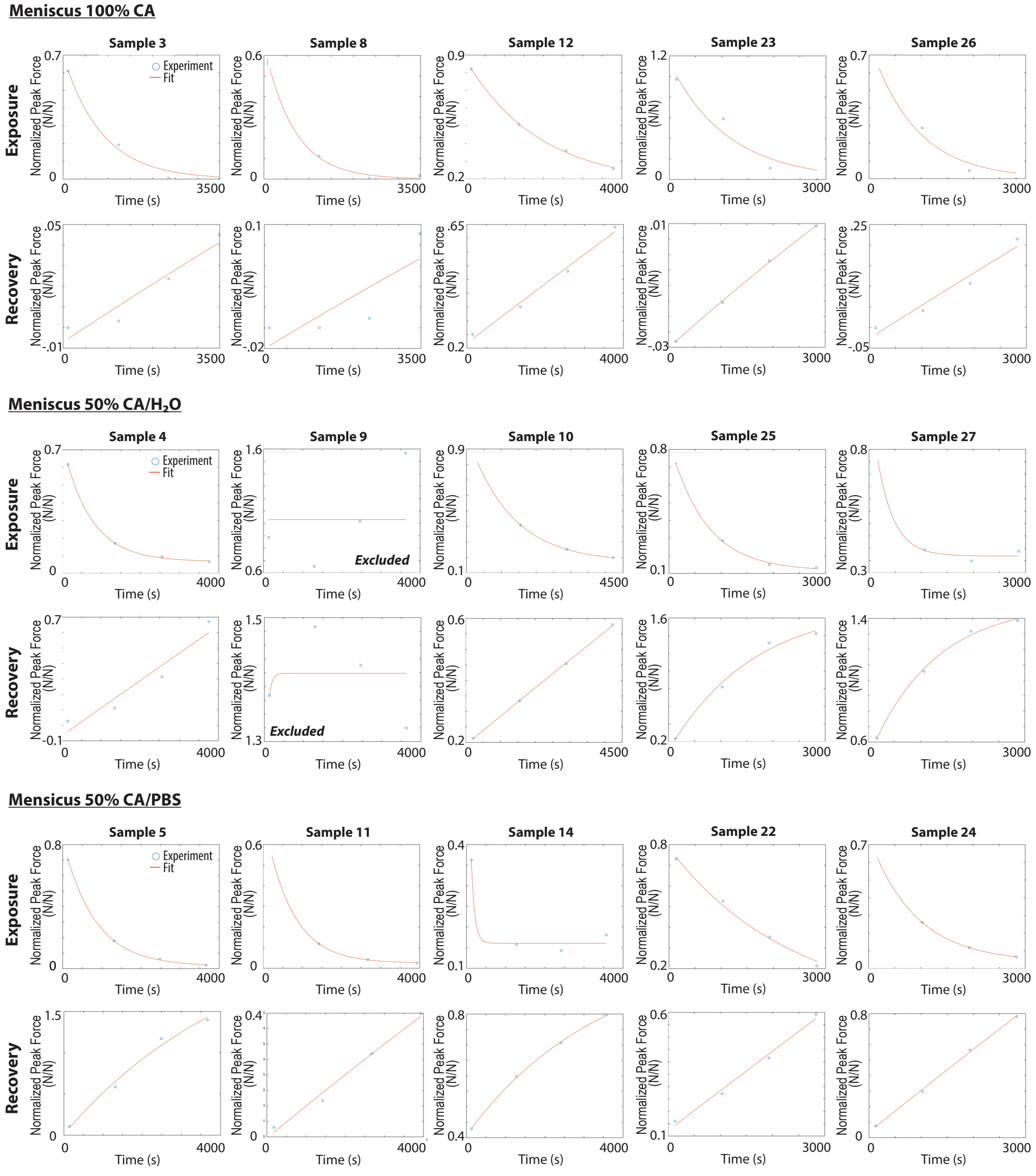


**Figure S4: Individual meniscus contrast agent and recovery data fits.** Blue markers represent raw data (normalized peak force), red line represents data fit. A monoexponential data fit was used for all contrast agent and recovery groups. Sample 9 (100% CA/$H_2O$) was excluded due to inconsistencies in the mechanical behavior.

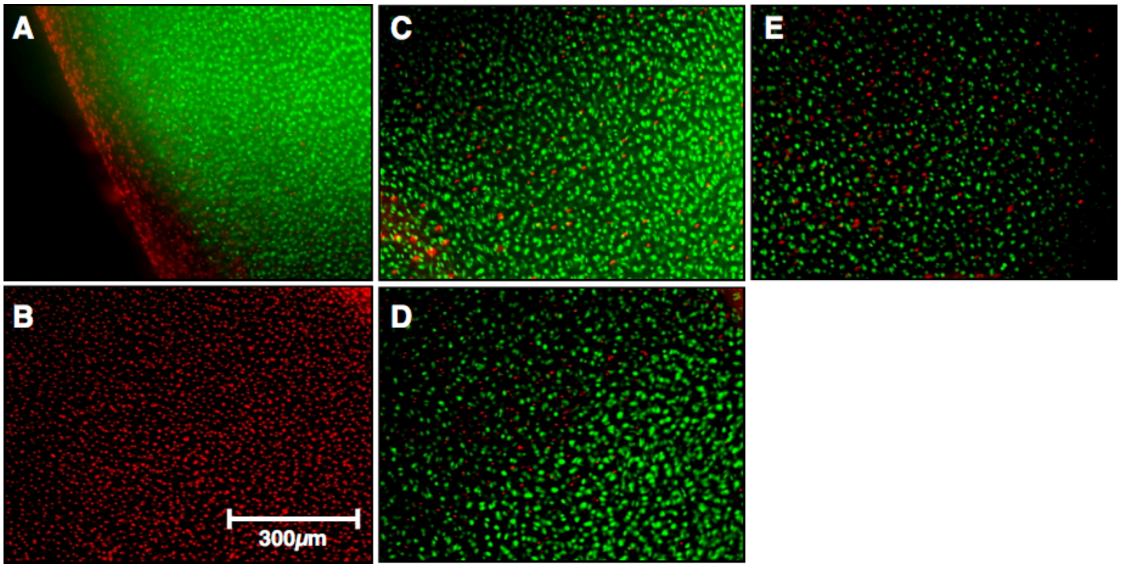


**Figure S5: Cell viability in cartilage explants was not affected by exposure to contrast agent dilutions.** As an initial assessment of possible toxic effects of contrast agent solutions on cartilage explants, cell viability was assessed with a live/dead cell staining kit (Invitrogen/Molecular Probes, Eugene, OR) after incubating the samples in their assigned contrast agent solution for 1 hour at 37˚C. Live/dead staining was visualized with a Zeiss Axiovert 200M Fluorescence Microscope (Göttingen, Germany). (A) PBS and (B) ethanol were used as negative and positive controls for cell death, respectively. Abundant live cells remained following exposure to (C) 100% CA, (D) 50% CA/$H_2O$, or (E) 50% CA/PBS, with no evidence of widespread cell death. A similar examination of MFC was not possible due to fluorescent imaging artifact caused by the abundant collagen network in MFC.

**Table S1.** A PubMed search was performed to identify all articles published in 2015-2018 involving CT arthrography of human subjects. The information tabulated is from the subset of articles reporting both the contrast agent used and information about its dilution. Estimated osmolalities for diluted contrast agents and ionic strengths were calculated based on reported properties of the contrast agents and of normal saline.

***Non-Ionic Contrast Agents***

| Author | Year | PMID | Agent | Dilution (% Full Strength) | Osmolality (mOsm/kg) | Est. Ionic Strength (mOsm/kg) |
|---|---|---|---|---|---|---|
| Abou Arab et. al.[1] | 2018 | 29713765 | Visipaque 270 | 100% | 290 | 0 |
| Abraham et. al.[2] | 2017 | 27923602 | Isovue 300 | 33% | 390 | 110 |
| Davis et. al.[3] | 2014 | 24491624 | Xenetix 300 | 100% | 695 | 0 |
| De Filippo et. al.[4] | 2016 | 27467868 | Iomeron 300 | 100% | 521 | 0 |
| Fox et. al.[5] | 2016 | 27144708 | Omnipaque 300 | 100% | 672 | 0 |
| Gondim Teixeira et. al.[6] | 2016 | 26700346 | Visipaque 270 | 100% | 290 | 0 |
| Guermazi et. al.[7] | 2014 | 24856241 | Omnipaque 300 | 35% | 410 | 110 |
| Henak et. al.[8] | 2014 | 25070373 | Isovue 300 | 33% | 390 | 110 |
| Kieffer et. al.[9] | 2014 | 24008621 | Visipaque 270 | 100% | 290 | 0 |
| Kirschke et. al.[10] | 2016 | 27891511 | Ultravist 300 | 67% | 480 | 60 |
| Knight et. al.[11] | 2017 | 28370312 | Isovue 300 | 33% | 390 | 110 |
| Magee, T.[12] | 2018 | 29549380 | Isovue 300 | 50% | 440 | 90 |
| Polat et. al.[13] | 2018 | 30371616 | Omnipaque 300 | 35% | 410 | 100 |
| Rauch et. al.[14] | 2018 | 30251928 | Visipaque 270 | 100% | 290 | 0 |
| Sahin et. al.[15] | 2014 | 25352941 | Iomeron 300 | 50% | 400 | 90 |
| Simeone et. al.[16] | 2017 | 28361351 | Isovue – M – 300 | 77% | 530 | 45 |
| Song et. al.[17] | 2018 | 29713219 | Ultravist 370 | 65% | 560 | 70 |
| Strobel et. al.[18] | 2014 | 24177809 | Iopamiro 300 | 50% | 440 | 90 |
| Suojarvi et. al.[19] | 2017 | 27456020 | Omnipaque 240 | 50% | 440 | 90 |
| Tobalem et. al.[20] | 2014 | 25415733 | Accupaque 300 | 75% | 530 | 50 |
| Wang et. al.[21] | 2018 | 29724673 | Omnipaque 300 | 70% | 530 | 60 |
| Zhang et. al.[22] | 2016 | 28002370 | Omnipaque 300 | 70% | 530 | 60 |

*Ionic Contrast Agents*

| Author | Year | PMID | Agent | Dilution (% Full Strength) | Osmolality (mOsm/kg) | Est. Ionic Strength (mOsm/kg) |
|---|---|---|---|---|---|---|
| Ahn et. al.[23] | 2014 | 25469085 | Telebrix 30 | 65% | 1200 | 590 |
| Cerny et. al.[24] | 2017 | 28409175 | Hexabrix | 100% | 600 | 300 |
| Choi et. al.[25] | 2016 | 26131912 | Telebrix 30 | 65% | 1200 | 590 |
| Ha et. al.[26] | 2017 | 28169108 | Telebrix 30 | 65% | 1200 | 590 |
| Ji et. al.[27] | 2014 | 24280955 | Telebrix 30 | 65% | 1200 | 590 |
| Jung et. al.[28] | 2018 | 28624855 | Telebrix 30 | 65% | 1200 | 590 |
| Kim et. al.[29] | 2014 | 24699852 | Telebrix 30 | 65% | 1200 | 590 |
| Kim et. al.[30] | 2016 | 25274092 | Telebrix 30 | 65% | 1200 | 590 |
| Kokkonen et. al.[31] | 2014 | 24249683 | Hexabrix 320 | 100% | 600 | 300 |
| Lyu et. al.[32] | 2014 | 27536568 | Telebrix 30 | 65% | 1200 | 590 |
| Moritomo et. al.[33] | 2015 | 25542437 | Urografin 60% | 50% | 810 | 405 |
| Myller et. al.[34] | 2017 | 27646147 | Hexabrix 320 | 50% | 440 | 220 |
| Nashikkar et. al.[35] | 2018 | 30174813 | Telebrix 30 | 65% | 1200 | 590 |
| Odri et. al.[36] | 2014 | 24797043 | Hexabrix 320 | 100% | 600 | 300 |
| Omoumi et. al.[37] | 2017 | 28189216 | Hexabrix 320 | 100% | 600 | 300 |
| Omoumi et. al.[38] | 2015 | 25450850 | Hexabrix 320 | 100% | 600 | 300 |
| Omoumi et. al.[39] | 2015 | 25377772 | Hexabrix 320 | 20% | 600 | 300 |
| Park et. al.[40] | 2015 | 25492036 | Telebrix 30 | 60% | 1100 | 180 |
| Park et. al.[41] | 2018 | 30021072 | Telebrix 30 | 60% | 1100 | 550 |
| Tabrizi et. al.[42] | 2015 | 25051918 | Urografin 60% | 100% | 1500 | 550 |
| Thienpont et. al.[43] | 2014 | 25382362 | Hexabrix 320 | 100% | 600 | 300 |
| van Tiel et. al.[44] | 2016 | 26851449 | Hexabrix 320 | 30% | 380 | 190 |
| Yoo et. al.[45] | 2017 | 28004173 | Telebrix 30 Meglumine | 65% | 1200 | 590 |
| Yoo et. al.[46] | 2017 | 27876489 | Telebrix 30 Meglumine | 65% | 1200 | 590 |

**Citations**


1. Abou Arab W, Rauch A, Chawki MB, et al. Scapholunate instability: improved detection with semi-automated kinematic CT analysis during stress maneuvers. *Eur Radiol*. 2018;28(10):4397-4406. doi:10.1007/s00330-018-5430-2.
2. Abraham CL, Knight SJ, Peters CL, Weiss JA, Anderson AE. Patient-specific chondrolabral contact mechanics in patients with acetabular dysplasia following treatment with peri-acetabular osteotomy. *Osteoarthr Cartil*. 2017;25(5):676-684. doi:10.1016/j.joca.2016.11.016.
3. Davis P, Stewart JJ, Hoover NG, Matthews BJ, Pahl DW, Bojescul JA. Use of CT-arthrography and ultrasound in ACL surgery during Operation Enduring Freedom in Afghanistan: a case report and practice recommendations. *Mil Med*. 2014;179(2):e240-4. doi:10.7205/MILMED-D-13-00248.
4. De Filippo M, Azzali E, Pesce A, et al. CT arthrography for evaluation of autologous chondrocyte and chondral-inductor scaffold implantation in the osteochondral lesions of the talus. *Acta Biomed*. 2016;87 Suppl 3:51-56.
5. Fox MG, Graham JA, Skelton BW, et al. Prospective Evaluation of Agreement and Accuracy in the Diagnosis of Meniscal Tears: MR Arthrography a Short Time After Injection Versus CT Arthrography After a Moderate Delay. *AJR Am J Roentgenol*. 2016;207(1):142-149. doi:10.2214/AJR.15.14517.
6. Gondim Teixeira PA, De Verbizier J, Aptel S, et al. Posterior Radioscaphoid Angle as a Predictor of Wrist Degenerative Joint Disease in Patients With Scapholunate Ligament Tears. *AJR Am J Roentgenol*. 2016;206(1):144-150. doi:10.2214/AJR.15.14606.
7. Guermazi A, Jomaah N, Hayashi D, et al. MR arthrography of the shoulder: optimizing pulse sequence protocols for the evaluation of cartilage and labrum. *Eur J Radiol*. 2014;83(8):1421-1428. doi:10.1016/j.ejrad.2014.04.030.
8. Henak CR, Abraham CL, Peters CL, Sanders RK, Weiss JA, Anderson AE. Computed tomography arthrography with traction in the human hip for three-dimensional reconstruction of cartilage and the acetabular labrum. *Clin Radiol*. 2014;69(10):e381-91. doi:10.1016/j.crad.2014.06.009.
9. Kieffer EM, Bouchaib J, Bierry G, Clavert P. CT arthrography and anatomical correlation of the bare area of the ulnar trochlear fossa: a risk of misdiagnosis of cartilage ulcerations. *Surg Radiol Anat*. 2014;36(5):481-486. doi:10.1007/s00276-013-1200-7.
10. Kirschke JS, Braun S, Baum T, et al. Diagnostic Value of CT Arthrography for Evaluation of Osteochondral Lesions at the Ankle. *Biomed Res Int*. 2016;2016:1-11. doi:10.1155/2016/3594253.
11. Knight SJ, Abraham CL, Peters CL, Weiss JA, Anderson AE. Changes in chondrolabral mechanics, coverage, and congruency following peri-acetabular osteotomy for treatment of acetabular retroversion: A patient-specific finite element study. *J Orthop Res*. 2017;35(11):2567-2576. doi:10.1002/jor.23566.
12. Magee T. Imaging of the post-operative shoulder: does injection of iodinated contrast in addition to MR contrast during arthrography improve diagnostic accuracy and patient throughput? *Skelet Radiol*. 2018;47(9):1253-1261. doi:10.1007/s00256-018-2927-3.
13. Polat G, Ogul H, Yalcin A, et al. Efficacy of the Rotational Traction Method in the Assessment of Glenohumeral Cartilage Surface Area in Computed Tomography Arthrography. *J Comput Assist Tomogr*. 2018. doi:10.1097/RCT.0000000000000809.
14. Rauch A, Arab WA, Dap F, Dautel G, Blum A, Gondim Teixeira PA. Four-dimensional

CT Analysis of Wrist Kinematics during Radioulnar Deviation. *Radiology*. 2018;289(3):750-758. doi:10.1148/radiol.2018180640.

15. Sahin M, Calisir C, Omeroglu H, Inan U, Mutlu F, Kaya T. Evaluation of Labral Pathology and Hip Articular Cartilage in Patients with Femoroacetabular Impingement (FAI): Comparison of Multidetector CT Arthrography and MR Arthrography. *Pol J Radiol*. 2014;79:374-380. doi:10.12659/PJR.890910.
16. Simeone FJ, Gill CM, Taneja AK, Torriani M, Bredella MA. Glenohumeral position during CT arthrography with arthroscopic correlation: optimization of diagnostic yield. *Skelet Radiol*. 2017;46(6):769-776. doi:10.1007/s00256-017-2613-x.
17. Song Y, Lee S, Lee BG, Joo YB, Song SY. The Diagnostic Reproducibility of Tomosynthesis for the Correlation between Acromiohumeral Distance and Rotator Cuff Size or Type. *Korean J Radiol*. 2018;19(3):417-424. doi:10.3348/kjr.2018.19.3.417.
18. Strobel K, Steurer-Dober I, Da Silva AJ, et al. Feasibility and preliminary results of SPECT/CT arthrography of the wrist in comparison with MR arthrography in patients with suspected ulnocarpal impaction. *Eur J Nucl Med Mol Imaging*. 2014;41(3):548-555. doi:10.1007/s00259-013-2584-7.
19. Suojarvi N, Haapamaki V, Lindfors N, Koskinen SK. Radiocarpal Injuries: Cone Beam Computed Tomography Arthrography, Magnetic Resonance Arthrography, and Arthroscopic Correlation among 21 Patients. *Scand J Surg*. 2017;106(2):173-179. doi:10.1177/1457496916659226.
20. Tobalem F, Dugert E, Verdun FR, et al. MDCT arthrography of the hip: value of the adaptive statistical iterative reconstruction technique and potential for radiation dose reduction. *AJR Am J Roentgenol*. 2014;203(6):W665-73. doi:10.2214/AJR.14.12821.
21. Wang J, Shao X, Huang M, Xin H, Zhang Z, Wang K. Predictors of Pain and Discomfort Associated with CT Arthrography of the Shoulder. *Acad Radiol*. 2018;25(12):1603-1608. doi:10.1016/j.acra.2018.04.003.
22. Zhang Q, Shi LL, Ravella KC, et al. Distinct Proximal Humeral Geometry in Chinese Population and Clinical Relevance. *J Bone Jt Surg Am*. 2016;98(24):2071-2081. doi:10.2106/JBJS.15.01232.
23. Ahn SJ, Hong SH, Chai JW, et al. Comparison of image quality of shoulder CT arthrography conducted using 120 kVp and 140 kVp protocols. *Korean J Radiol*. 2014;15(6):739-745. doi:10.3348/kjr.2014.15.6.739.
24. Cerny M, Omoumi P, Larbi A, et al. CT arthrography of adhesive capsulitis of the shoulder: Are MR signs applicable? *Eur J Radiol Open*. 2017;4:40-44. doi:10.1016/j.ejro.2017.03.002.
25. Choi BH, Kim NR, Moon SG, Park JY, Choi JW. Superior Labral Cleft after Superior Labral Anterior-to-Posterior Tear Repair: CT Arthrographic Features and Correlation with Clinical Outcome. *Radiology*. 2016;278(2):441-448. doi:10.1148/radiol.2015142431.
26. Ha YC, Lee YK, Koo KH, Kwon KB, Song SH. Prevalence and clinical significance of hypertrophic labrum in non-dysplastic hips. *J Orthop Sci*. 2017;22(3):512-516. doi:10.1016/j.jos.2017.01.010.
27. Ji HM, Baek JH, Kim KW, Yoon JW, Ha YC. Herniation pits as a radiographic indicator of pincer-type femoroacetabular impingement in symptomatic patients. *Knee Surg Sport Traumatol Arthrosc*. 2014;22(4):860-866. doi:10.1007/s00167-013-2777-4.
28. Jung HG, Kim NR, Jeon JY, et al. CT arthrography visualizes tissue growth of osteochondral defects of the talus after microfracture. *Knee Surg Sport Traumatol*

*Arthrosc*. 2018;26(7):2123-2130. doi:10.1007/s00167-017-4610-y.
29. Kim JY, Chung SW, Kwak JY. Morphological Characteristics of the Repaired Labrum According to Glenoid Location and Its Clinical Relevance After Arthroscopic Bankart Repair: Postoperative Evaluation With Computed Tomography Arthrography. *Am J Sport Med*. 2014;42(6):1304-1314. doi:10.1177/0363546514528791.
30. Kim SH, Choi JY, Yoo HJ, Hong SH. External rotation and active supination CT arthrography for the postoperative evaluation of type II superior labral anterior to posterior lesions. *Knee Surg Sport Traumatol Arthrosc*. 2016;24(1):134-140. doi:10.1007/s00167-014-3350-5.
31. Kokkonen HT, Suomalainen JS, Joukainen A, et al. In vivo diagnostics of human knee cartilage lesions using delayed CBCT arthrography. *J Orthop Res*. 2014;32(3):403-412. doi:10.1002/jor.22521.
32. Lyu SH, Kwak YH, Lee YK, Ha YC, Koo KH. Correlation of Structural Bony Abnormalities and Mechanical Symptoms of Hip Joints. *Hip Pelvis*. 2014;26(2):115-123. doi:10.5371/hp.2014.26.2.115.
33. Moritomo H, Arimitsu S, Kubo N, Masatomi T, Yukioka M. Computed tomography arthrography using a radial plane view for the detection of triangular fibrocartilage complex foveal tears. *J Hand Surg Am*. 2015;40(2):245-251. doi:10.1016/j.jhsa.2014.10.051.
34. Myller KA, Turunen MJ, Honkanen JT, et al. In Vivo Contrast-Enhanced Cone Beam CT Provides Quantitative Information on Articular Cartilage and Subchondral Bone. *Ann Biomed Eng*. 2017;45(3):811-818. doi:10.1007/s10439-016-1730-3.
35. Nashikkar PS, Rhee SM, Desai C V, Oh JH. Is Anatomical Healing Essential for Better Clinical Outcome in Type II SLAP Repair? Clinico-Radiological Outcome after Type II SLAP Repair. *Clin Orthop Surg*. 2018;10(3):358-367. doi:10.4055/cios.2018.10.3.358.
36. Odri GA, Frioux R, Redon H, et al. Reliability of a new hip lateral view to quantify alpha angle in femoroacetabular impingement. *Orthop Traumatol Surg Res*. 2014;100(4):363-367. doi:10.1016/j.otsr.2014.01.011.
37. Omoumi P, Michoux N, Larbi A, et al. Multirater agreement for grading the femoral and tibial cartilage surface lesions at CT arthrography and analysis of causes of disagreement. *Eur J Radiol*. 2017;88:95-101. doi:10.1016/j.ejrad.2016.12.026.
38. Omoumi P, Michoux N, Roemer FW, Thienpont E, Vande Berg BC. Cartilage thickness at the posterior medial femoral condyle is increased in femorotibial knee osteoarthritis: a cross-sectional CT arthrography study (Part 2). *Osteoarthr Cartil*. 2015;23(2):224-231. doi:10.1016/j.joca.2014.08.017.
39. Omoumi P, Rubini A, Dubuc JE, Vande Berg BC, Lecouvet FE. Diagnostic performance of CT-arthrography and 1.5T MR-arthrography for the assessment of glenohumeral joint cartilage: a comparative study with arthroscopic correlation. *Eur Radiol*. 2015;25(4):961-969. doi:10.1007/s00330-014-3469-2.
40. Park JY, Chung SW, Kumar G, et al. Factors affecting capsular volume changes and association with outcomes after Bankart repair and capsular shift. *Am J Sport Med*. 2015;43(2):428-438. doi:10.1177/0363546514559825.
41. Park JY, Lee JH, Chung SW, Oh KS, Noh YM, Kim SJ. Does Anchor Placement on the Glenoid Affect Functional Outcome After Arthroscopic Bankart Repair? *Am J Sport Med*. 2018;46(10):2466-2471. doi:10.1177/0363546518785968.
42. Tabrizi PR, Zoroofi RA, Yokota F, Tamura S, Nishii T, Sato Y. Acetabular cartilage

segmentation in CT arthrography based on a bone-normalized probabilistic atlas. *Int J Comput Assist Radiol Surg*. 2015;10(4):433-446. doi:10.1007/s11548-014-1101-1.

43. Thienpont E, Schwab P-E, Omoumi P. Wear patterns in anteromedial osteoarthritis of the knee evaluated with CT-arthrography. *Knee*. 2014;21:S15-S19. doi:10.1016/s0968-0160(14)50004-x.
44. van Tiel J, Siebelt M, Reijman M, et al. Quantitative in vivo CT arthrography of the human osteoarthritic knee to estimate cartilage sulphated glycosaminoglycan content: correlation with ex-vivo reference standards. *Osteoarthr Cartil*. 2016;24(6):1012-1020. doi:10.1016/j.joca.2016.01.137.
45. Yoo JI, Ha YC, Lee HJ, Lee JY, Lee YK, Koo KH. No difference in prevalence of radiographic subspinal impingement of the hip between symptomatic and asymptomatic subjects. *Knee Surg Sport Traumatol Arthrosc*. 2017;25(6):1951-1957. doi:10.1007/s00167-016-4402-9.
46. Yoo JI, Ha YC, Lee YK, Lee GY, Yoo MJ, Koo KH. Morphologic Changes and Outcomes After Arthroscopic Acetabular Labral Repair Evaluated Using Postoperative Computed Tomography Arthrography. *Arthroscopy*. 2017;33(2):337-345. doi:10.1016/j.arthro.2016.08.022.
